# Scuticociliate Detection and Microbiome Composition in Museum Collections of *Diadema Antillarum*

**DOI:** 10.1101/2025.11.24.690235

**Authors:** Brayan Vilanova-Cuevas, Ian Hewson

**Affiliations:** Cornell Oceans and Department of Microbiology, Cornell University, Ithaca, New York, USA

## Abstract

**Background:** The mass mortality of the long-spined sea urchin Diadema antillarum has caused widespread ecological changes across Caribbean reefs, with recent studies identifying the etiological agent as pathogenic ciliate designated as a D. antillarum Scuticociliatosis Philaster-clade (DaScPc). The origin and ecological trajectory of DaScPc remain unresolved, raising critical questions about whether it represents a novel introduction or a resident commensal symbiont that transitioned into pathogenicity.

**Methods:** To address this, we tested 50 individual preserved museum specimens of D. antillarum collected between 1960 and 2020, with targeted PCR amplification of ciliate 18S, 28S, and 5.8S/ITS rRNA genes for spine, body wall, and coelomic fluid samples (n=100). Following up on recent work that identified bacterial biomarkers of DaSc, we also characterized the microbial communities associated with these museum specimens using 16S rRNA amplicon sequencing. Results. Our results reveal the presence of identical DaScPc 18S rRNA sequences in 21% of tested samples, 28S rRNA PCR yielded sequences at 96-98 % nt identity in only 2% of the tested samples, and we got no amplification from the 5.8S/ITS region. While these findings suggest possible long-term persistence or repeated emergence of this ciliate, the lack of 28S rRNA matches and lack of detection of ITS2 demonstrates that DaScPc 18S rRNA gene detections may be false positives for the ciliate over a highly conserved rRNA region. The microbial composition of the samples didn’t yield any of the previously identified disease-associated bacterial biomarkers and showed large shifts in the overall microbial community based on collection period and the facility where the samples are housed. This study demonstrates that museum-preserved echinoderm tissues retain ecologically informative microbial DNA and establishes a molecular framework for disentangling pathogen provenance and its caveats. It also highlights the value and limitations of natural history collections in reconstructing marine disease ecology.

## Introduction

The long-spined sea urchin *Diadema antillarum* was a keystone herbivore on Caribbean coral reefs (Lessios, 2016), playing a vital role in maintaining reef health by controlling macroalgal density. *D. antillarum* experienced a catastrophic mass mortality event in the Caribbean in 1983–1984, with ∼ 95% mortality at affected sites contributing to increased macroalgal density and coral decline (Lessios, Robertson & Cubit, 1984). The causative agent of the mass mortality was never established, and no specimens of affected urchins remain today. Another localized mass mortality affected *D. antillarum* in the Florida Keys in early 1991 (Forcucci, 1994), and in 2022 this species was decimated at sites across the eastern Caribbean (Hylkema et al., 2023). The 2022 mass mortality event was caused by the *D. antillarum* scuticociliatosis Philaster-clade (DaScPc) (Hewson et al., 2023). Subsequent mass mortalities of urchins within the Diadematidae family were reported in the Mediterranean, Red Sea, and the Indian Ocean and molecular diagnostic using 18S rRNA confirmed DaScPc in affected urchins from these locations (Ritchie et al., 2024; Roth et al., 2024; Quod et al., 2025). Recent studies have demonstrated that DaScPc, detected by 18S rRNA sequencing, may persist on surfaces of coral reefs after mass mortality, including on prominent coral species and other abiotic surfaces (e.g. boat hulls) (Vilanova-Cuevas et al., 2025b). Attempts to PCR amplify DaScPc in available Caribbean coral and coral reef plankton microbial DNA extracts collected prior to the mortality event were unsuccessful (Vilanova-Cuevas et al., 2025b). These results inspired the hypothesis that DaScPc may have been a novel introduction to the affected region in 2022. However, it is also possible that it was present in urchins prior to 2022, since previous surveys were opportunistic and did not focus on urchins.

One avenue for investigating historical disease dynamics is through the study of historic DNA (hDNA), which refers to genetic material extracted from preserved museum specimens (Arning & Wilson, 2020). hDNA can be highly degraded due to chemical interactions with fixatives but provides an invaluable resource for retrospective genomic studies (Raxworthy & Smith, 2021). Scientific (museum) collections serve as critical repositories for studying historical biodiversity and ecological changes over time. These collections allow researchers to test hypotheses about disease emergence and persistence by detecting pathogens in preserved specimens (Raxworthy & Smith, 2021). If DaScPc DNA can be identified in *D. antillarum* specimens collected prior to 2022, it may indicate that it associates with *D. antillarum* as a normal constituent of the holobiont that could be environmentally triggered rather than novelly introduced.

The use of preserved museum specimens also allows researchers to study variation in holobiont microbiome composition (Arning & Wilson, 2020). Recent work has described microbiome composition of century old, preserved specimens and these showed no correlation between the microbiome and collection time, suggesting preservation of material is sufficient to retain their composition at the time of collection (Chalifour, Elder & Li, 2022). Studies have also investigated host-microbe interactions, in fish, by examining symbiont presence or absence (Gould, Fritts-Penniman & Gaisiner, 2021). Echinoids depend on a core microbial community for nutrient acquisition, digestion and overall health, where diseases impact their microbiomes (Shaw et al., 2023; Park et al., 2023; Bengtsson et al., 2025). Most recently, a bacterial biomarker, *Fangia hongkongensis*, was associated with DaSc infected urchin specimens collected from the Caribbean and the Western Indian Ocean (Vilanova-Cuevas et al., 2025a). This finding provides a possible bacterial biomarker for scuticociliatosis in Diadematidae urchins that serves as a potential target for retrospective identification of DaScPc in museum specimens.

We examined preserved *D. antillarum* specimens from the US National Museum of Natural History at the Smithsonian Institution and the Marine Invertebrate Collection of the Florida Marine and Wildlife Conservation Commission, spanning five decades (1965–2015), for DaScPc and other microorganisms. We interrogated specimens by PCR targeting the 18S and 28S rRNA genes and intergenic spacer 2 (ITS2) of DaScPc. Furthermore, we examined microbiome composition via 16S rRNA amplicon sequencing to elucidate whether microbial markers of disease were present in congruence with identification of DaScPc. This research highlights the important role of museum-preserved specimens in disease ecology.

## Materials and Methods

### Museum Sample Collection and Processing

Samples of *D. antillarum* spine and coelomic fluid were obtained from the National Museum of Natural History at the Smithsonian Institution and combined spine/body wall samples and coelomic fluid were collected from the Marine Invertebrate Collection of the Florida Marine and Wildlife Conservation Commission. A total of 100 samples were taken from 50 individual urchin specimens, spanning collection years from 1965 to 2015. Specimens were mostly preserved with multiple urchins per jar (n = 28), but some were preserved as single urchins in jars (n = 22). The exact preservation conditions upon collection are not documented. Marine invertebrates are frequently fixed with formalin or other fixatives (which inhibit PCR amplification), and it is unclear how long after collection from the wild that specimens were fixed or preserved, which may affect microbiome composition and the proliferation of ciliates, since these frequently participate in echinoderm tissue breakdown as saprobes (Hewson et al., 2023). All samples had been maintained in 75–90% ethanol at their respective institutions. Detailed information regarding collection vessels, location, and museum specimen IDs is provided in Supplementary Table 1.

Coelomic fluid sample were collected using a sterile syringe and 25G needle inserted through the echinoid peristomal membrane (0.05 – 0.1 mL per specimen). Several specimens did not have intact tests or peristomal membranes at the time of observation and so coelomic fluid was not sampled. A spine from each specimen was removed by applying gentle pressure using clean forceps. For specimens obtained from the Florida Marine and Wildlife Conservation Commission, a piece of spine and body wall was collected by carefully breaking open the test with sterile forceps. All samples were immediately placed into 1 mL of RNA Later (Invitrogen, Carlsbad, CA, USA) and transported to the laboratory at Cornell University at ambient temperature. The samples were kept at −80°C until DNA extraction was performed using the Quick-DNA Insect/Tissue kit (Zymo Research, Irvine, CA, USA) per the manufacturer’s protocols.

### 18S and 28S rRNA Gene Amplification and Read Processing

18S rRNA gene amplification: DNA extracts of museum samples were amplified using primer 384F (3’-YTBGATGGTAGTGTATTGGA-5’) and 1147 (3’-AACCTTGGAGACCTGAT-5’) for a product size of ∼750 bp of the 18S rRNA (Dopheide et al., 2008). A nested PCR was performed on the PCR product amplification using 634f-scutico (3’-TTGCAATGAGAACAACGTAA-5’) and the same reverse primers before, for a product ∼450 bp (Vilanova-Cuevas et al., 2023). Amplification for the 18S rRNA was carried out in 50 µl reactions containing 1X PCR Buffer (ThermoFisher), 2.5 mM MgCl, 0.2 mM PCR Nucleotide Mix (Promega), 10 µM of each primer, 5U Taq DNA polymerase (ThermoFisher), 0.02 ng/µL BSA, and 2 µL of template DNA. Negative controls for each run comprised 1 µL of nuclease free water, and positive controls comprised µL of DNA extracted from Culture FWC2. The thermal cycling conditions for the 28S rRNA included an initial denaturation at 95°C for 3 minutes, followed by 30 cycles of denaturation at 95°C for 30 seconds, annealing at 52°C for 90 seconds, and extension at 72°C for 120 seconds. The thermal cycling conditions for the 18S rRNA included an initial denaturation at 95°C for 3 minutes, followed by 30 cycles of denaturation at 95°C for 30 seconds, annealing at 54°C for 30 seconds, and extension at 72°C for 30 seconds. A final extension step at 72°C for 5 minutes was performed to complete the reaction.

28S rRNA gene amplification: The primers were designed using Primer3 (Rozen & Skaletsky, 1999), based on a PCR amplified 28S rRNA from DaScPc culture FWC2 and related scuticociliates retrieved from NCBI (Supplementary Figure 1) (Rozen & Skaletsky, 1999; Hewson et al., 2023). DNA extracts from museum samples were amplified using primer 28F (3’-ACSCGCTGRAYTTAAGCAT-5’) and 28R (3’-AACCTTGGAGACCTGAT-5’) for a predicted product size of ∼1850 nt. Because initial amplification using 28F/28R resulted in low amplicon amount, amplicons were subject to nested PCR using primer Scutico-F28S (3’-CTGCGAAGGAAAGGTGAAAAGA-5’), designed from DaScPc culture FWC2 28S rRNA, and primer 28R, with a predicted product size of ∼1500 nt. Amplification for both 28S rRNA steps was carried out in 50 µl reactions containing 1X PCR Buffer (ThermoFisher), 2.5 mM MgCl, 0.2 mM PCR Nucleotide Mix (Promega), 10 µM of each primer, 5U Taq DNA polymerase (ThermoFisher), 0.02 ng µL^-1^ BSA, and 2 µl of template DNA. The thermal cycling conditions for the first 28S rRNA step included an initial denaturation at 95°C for 3 minutes, followed by 30 cycles of denaturation at 95°C for 30 seconds, annealing at 52°C for 90 seconds, and extension at 72°C for 120 seconds. A final extension step at 72°C for 5 minutes was performed to complete the reaction. The nested amplification followed the previously described thermal cycling conditions except that annealing temperature was lowered to 53°C for 90 seconds.

5.8S/ITS rRNA gene amplification: The primers were designed using Primer3, based on a PCR amplified 5.8S/ITS region from DaScPc culture FWC2 and related scuticociliates retrieved from NCBI OP896845.1) (Rozen & Skaletsky, 1999; Hewson et al., 2023). DNA extracts from museum samples were amplified using primer 5.8IT-F (3’-GTAGGTGAACCTTCGGAAGGATCATTA −5’) and 5.8ITS-R (3’-TACTGATATTGCTTAGTTCAGCGG −5’) for a predicted product size of ∼518 nt. Amplification for 5.8S rRNA steps was carried out in 50 µL reactions containing 1X PCR Buffer (ThermoFisher), 2.5 mM MgCl, 0.2 mM PCR Nucleotide Mix (Promega), 10 µM of each primer, 5U Taq DNA polymerase (ThermoFisher), 0.02 ng µl-1 BSA, and 2 µL of template DNA. The thermal cycling conditions for the first 5.8S rRNA step included an initial denaturation at 95°C for 3 minutes, followed by 30 cycles of denaturation at 95°C for 30 seconds, annealing at 58°C for 90 seconds, and extension at 72°C for 120 seconds. A final extension step at 72°C for 5 minutes was performed to complete the reaction.

PCR products, along with a standard ladder (Promega), were separated on 1% agarose gels in 1X TBE buffer by electrophoresis at 85V for 1 hour. Gels were subsequently stained with SYBR Gold (10X) and visualized using a BioRad ChemiDoc system. Amplicons that matched the expected size were purified using the Zymo Clean & Concentrator-5 kit and Sanger sequenced at the Cornell University Biotechnology Research Center. All sequences generated in this study have been deposited in GenBank under accession numbers PX138965 to PX139012.

### 16S rRNA Gene Amplification and Read Processing

Bacterial and archaeal communities in tissue samples (n = 51) were analyzed using dual-barcoded PCR amplification and sequencing of the V4 region of the 16S rRNA gene (Kozich et al., 2013). Each 40 µL PCR reaction contained 1× PCR master mix (One-Taq Quick-Load 2× Master Mix with Standard; New England Biolabs, Ipswich, MA, USA), 0.125 µM of each barcoded primer (515f; 5′-GTG YCA GCM GCC GCG GTA A3′ and 806r; 5′-GGA CTA CNV GGG TWT CTA AT-3′), and 2 µL of template. Thermocycling conditions followed Apprill et al. 2015. The 16S rRNA amplicons were pooled at equal concentrations using the SequalPrep Normalization Plate kit (Invitrogen) and sequenced on an Illumina MiSeq platform (2 × 250 paired end) at Cornell University’s Biotechnology Research Center. The 16S rRNA gene amplicon sequences were submitted to EBI (EBI Accession no. PRJEB97190 and ERP179787).

Raw sequence data were preprocessed in QIITA under Study ID 15818, where samples were demultiplexed and quality control retained samples with a Phred Score above 30 (Gonzalez et al., 2018). Sequences were trimmed at 250 nt, and Deblur was used to denoise the sequences, creating an initial amplicon sequencing variants (ASVs) table (Amir et al., 2017). Taxonomy was assigned with a 97% identity confidence threshold, using the Silva-138-99-515-806-nb-classifier (V.138), chloroplast and mitochondrial sequences were removed from the ASV table, and singletons and doubletons were also removed (Quast et al., 2013). To control for sampling depth differences across libraries, all samples were rarefied to an even depth of 3,000 sequences. Data were then exported to R for alpha and beta diversity calculations, as well as taxonomic composition analyses using Phyloseq (v.1.46.0) and a graphical summary created using ggplot2 (v.3.5.0) (McMurdie & Holmes, 2013; Wickham, 2016)

Differences in richness and diversity were assessed using the Chao1 and Shannon index, respectively (Shannon, 1948; Chao, 1984; Keylock, 2005). Significant differences in alpha diversity were determined using the Wilcoxon test with ggpubr (v.0.6.0) (Kassambara, 2023). Beta diversity analysis was performed using Bray-Curtis dissimilarities index and visualized with a PCoA, using Phyloseq and ggplot2 (Bray & Curtis, 1957; McMurdie & Holmes, 2013; Wickham, 2016). Taxonomic composition was assessed for genus level composition surpassing 5% relative abundance using Phyloseq and ggplot2 (McMurdie & Holmes, 2013; Wickham, 2016). The statistical comparison for the analysis of variance using distance matrix was done with a PERMANOVA test, using the Adonis function in vegan (v.2.6-4) (“vegan: Community Ecology Package,” 2022). To assess correlations between microbial community composition and metadata variables (decade, project, year, and DaScPc18S status), taxonomic counts were collapsed to the genus level and transformed to relative abundances. Metadata variables were encoded numerically, and Pearson correlation coefficients and associated p-values between each genus and each metadata variable were calculated (McMurdie & Holmes, 2013; Neuwirth, 2022; Wickham et al., 2023, 2024, 2025)(McMurdie & Holmes, 2013; Neuwirth, 2022; Wickham et al., 2023, 2024, 2025). Resulting correlations were visualized as a heatmap, with annotations displaying both correlation coefficients and significance levels (Kolde, 2019).

## Results

### PCR Detection of DaScPc Marker Genes

PCR amplification of the 18S, 28S, and 5.8S rRNAs was performed on 100 urchin DNA extracts (Table 3-1). PCR of 18S rRNA yielded amplicons in 32 DNA extracts, comprising 20 spine, spine/body wall, and 13 coelomic fluid samples. Phylogenetic analysis revealed that 21 (n = 16 individual *D. antillarum* specimens) of these sequences fell within a clade including DaScPc-infected urchins from the Caribbean in 2022, DaScPc-infected urchins from Reunion Island in 2023 and the DaScPc culture FWC2 (Figure 1A) (Roth et al., 2024). Of these 16 positive specimens, 4 were from museum jars containing multiple urchins, but other urchins within the same jar did not yield DaScPc amplicons. Five individual *D. antillarum* specimens yielded DaScPc sequences in both spine/body wall and coelomic fluid, while the remaining 11 specimens yielded DaScPc amplicons in either coelomic fluid or spine/body wall but not both. DaScPc-positive specimens were collected between 1965 and 2014. Two sequences, from different specimens and facilities, were most similar to Acropora/CHN/2009 (Accession: HM030718.1 and HM030719.1), initially recovered from corals in Hainan (Qiu et al., 2010), and recently described as associated with corals in the Caribbean (Vilanova-Cuevas et al., 2025b). The remaining sequences matched *Paralembus digitiformis*. Amplification of 28S rRNA was less successful, with only two DNA extracts (n = 2 individual specimens) from 1982 and 1983 yielding amplicons. Both sequences were most similar to DaScPc culture FWC2, but not identical across the locus (96-98 % nucleotide identity (Figure 2). Amplification of the 5.8S/ITS region didn’t yield any amplicon for any of the tested museum specimens.

**Figure 1.**
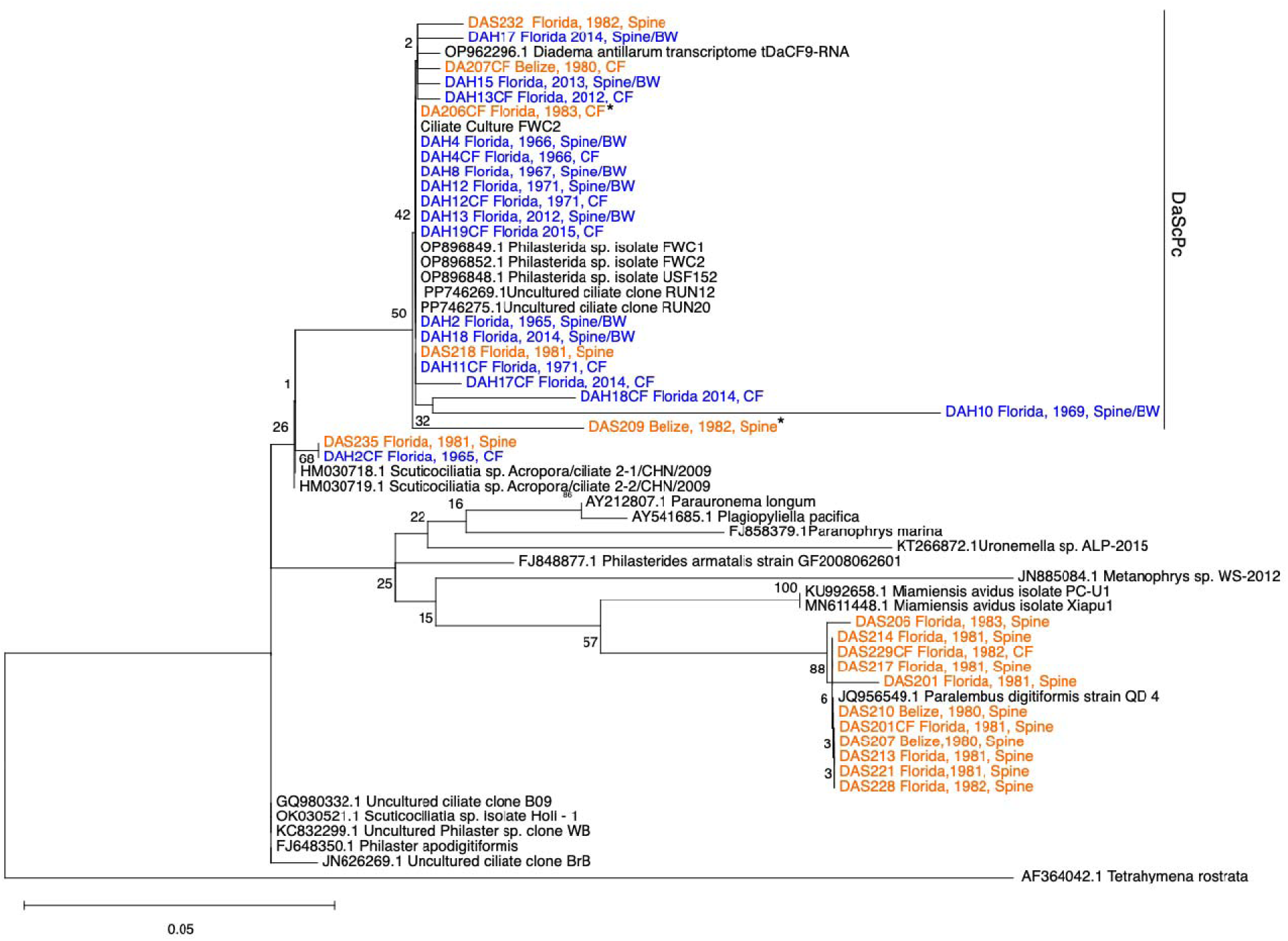
Phylogenetic analysis of 18S rRNA sequences obtained from US National Museum of Natural History (Smithsonian) Echinoderm Collection (Orange) and the University of South Florida Echinoderm Collection (Blue). The tree is based on 326 bp of the 18S, overlapping portion of each were aligned with MUSCLE (Edgar, 2004). The phylogenetic construction was based on Maximum Likelihood, following the Kimura 2-parameter model, with gamma rate nucleotide substitution method and the Nearest-Neighbor-Interchange heuristic model. Bootstrap values are based on 100 iterations of tree clustering.

**Figure 2.**
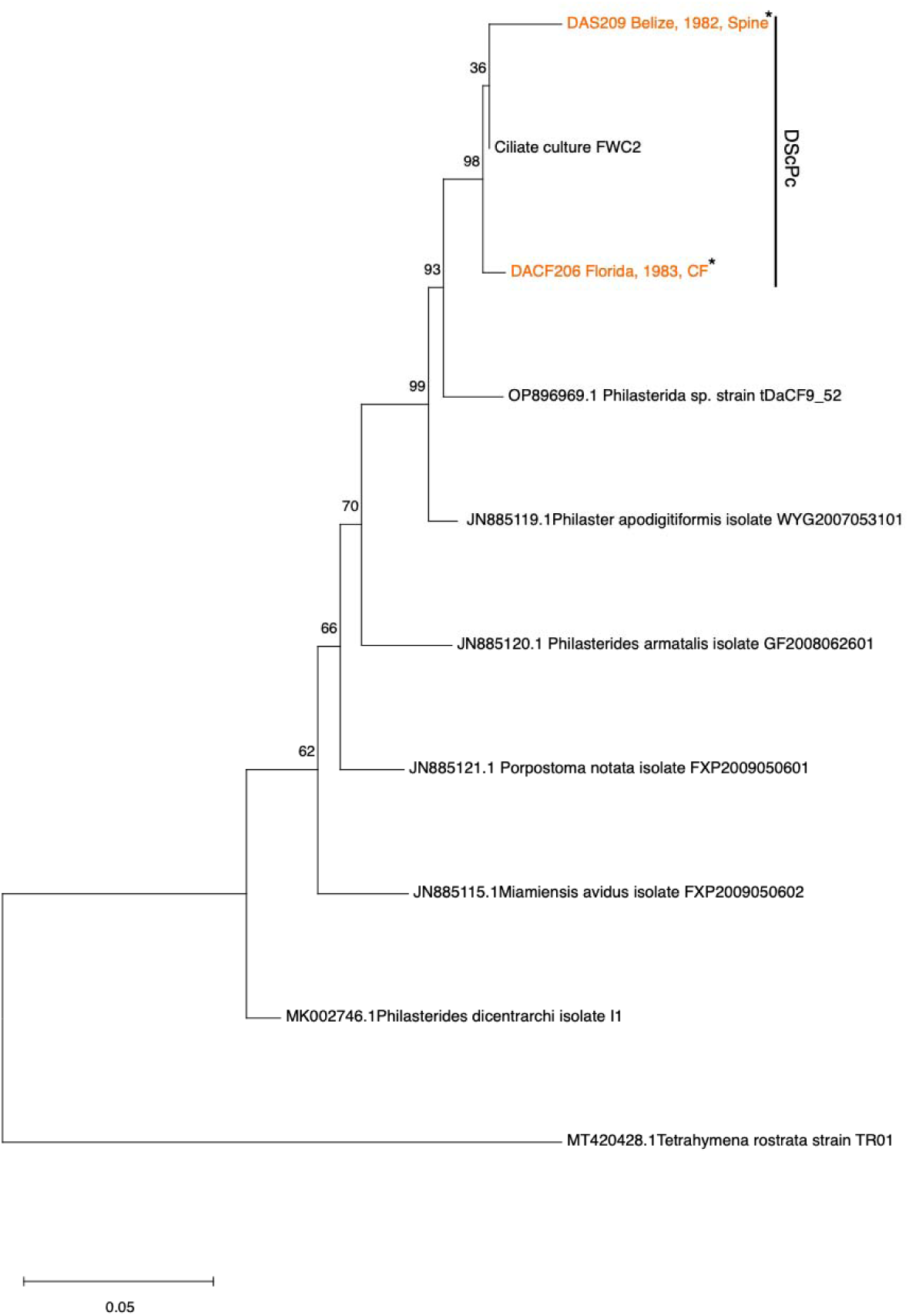
Phylogenetic analysis of 28S rRNA sequences obtained from US National Museum of Natural History (Smithsonian) Echinoderm Collection (Orange) and the University of South Florida Echinoderm Collection (Green). The tree is based on 1140 bp of the 28S rRNA, overlapping portion of each were aligned with MUSCLE (Edgar, 2004). The phylogenetic construction was based on Maximum Likelihood, following the Kimura 2-parameter model, with gamma rate nucleotide substitution for the and the Nearest-Neighbor-Interchange heuristic model. Bootstrap values are based on 100 iterations of tree clustering.

### Microbial Composition Analysis

Microbiome (16S rRNA gene) amplicon analysis was performed on 64 *D. antillarum* tissue samples which included spines (n=37), from the US National Museum of Natural History (Smithsonian) collection, and spine/body wall (n=13), and coelomic fluid (n=13) from the Florida Fish and Wildlife Conservation Commission. Alpha diversity (Chao1 and Shannon indices; Figure 3A), were variable across decades, with samples collected between 1960-1969 and 1980-1989 bearing significantly lower microbial richness (comparison by decade, Wilcoxon test comparison, p-val 0.04 and 8.6×10^-5^ respectively) and diversity (comparison by decade, Wilcoxon test comparison, p-val 0.02 and 6.7×10^-6^ respectively) when compared to samples collected between 2010-2019. Beta diversity revealed distinct clustering patterns (Principal Coordinates Analysis with Bray-Curtis dissimilarity metrics; Figure 3B), indicating significant differences in microbial composition across facilities and collection periods (PERMANOVA;R² = 0.31, p < 0.05 for collection decade and R² = 0.22, p < 0.05 for facility). There was no significant dissimilarity based on the collection location of the samples, which we divided into separate analysis by facility to avoid false dissimilarity related to facility differences previously shown (Supplementary Figure 2).

**Figure 3.**
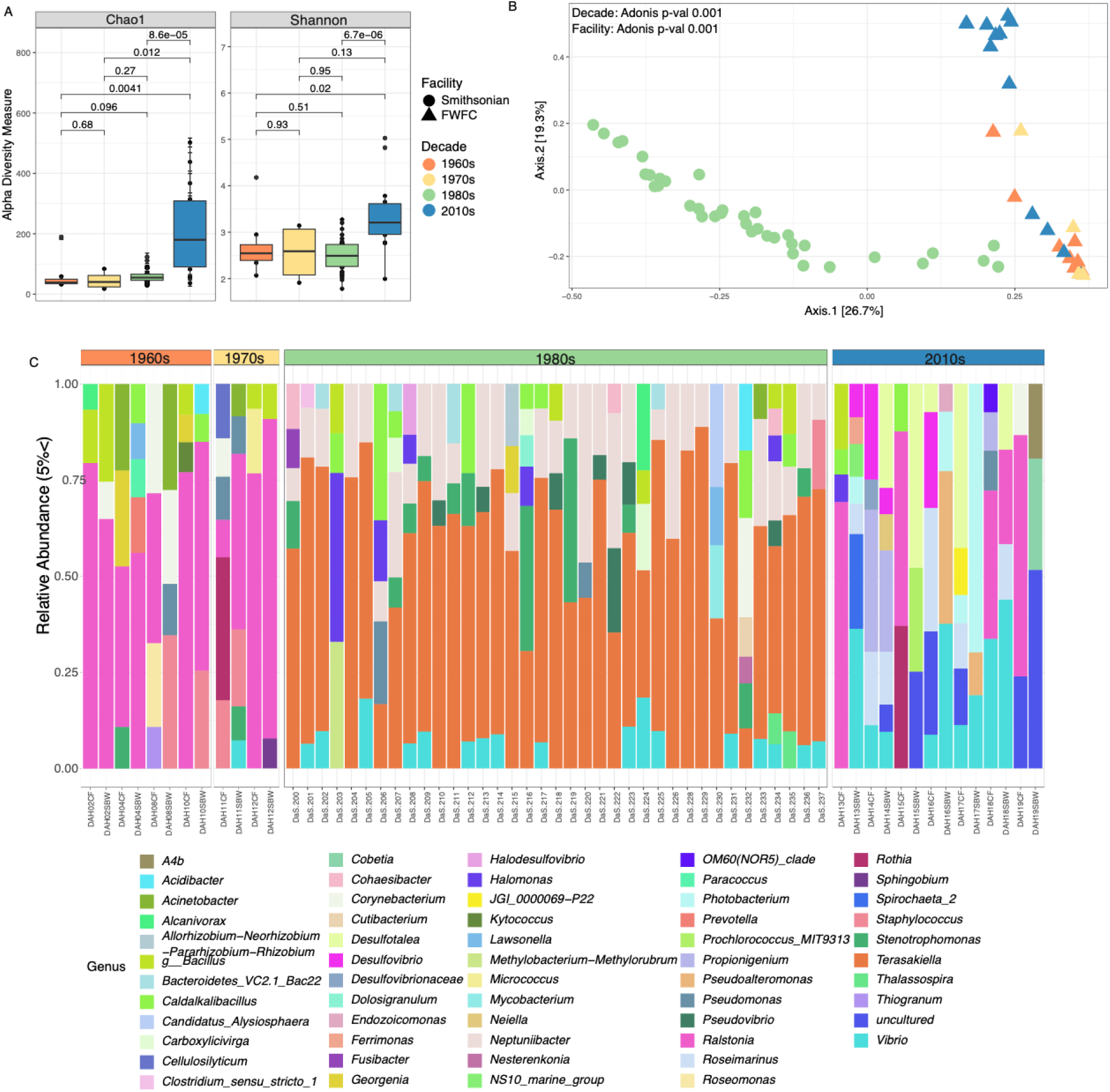
Microbial community analysis comparing historic samples based on the decade of acquisition and facility. Alpha diversity analysis (A) shows both richness (Chao1) and diversity (Shannon) across the four decades analyzed, p-values represent Wilcoxon test comparisons. Beta diversity analysis (B) was computed using Bray-Curtis dissimilarities and a PCoA visualization, with statistical comparison by variable measured using ADONIS. (C) Genus level taxonomic composition by samples, grouped by the decades, showing genera surpassing 5% relative abundance.

The relative abundance of bacterial genera varied between decades, reflecting shifts in microbial community composition influenced by both temporal changes and facility-specific factors (Figure 3C). Taxonomic composition in libraries prepared from specimens collected from 1960-1969, 1970-1979, and 2010-2019, which were obtained from the Marine Invertebrate Collection of the Florida Marine and Wildlife Conservation Commission, were broadly different from those 1980s samples collected from National Museum of Natural History at the Smithsonian Institution. Due to lack of specific location of collection, difference in facility, and time of collection, we are unable to distinguish specific factors contributing to microbial variability. In libraries prepared between 1960-1969, the amplicon libraries were primarily composed of *Ralstonia* (41%), followed by *Corynebacterium* (5.4%) and *Bacillus* (5.2%). In libraries prepared from specimens collected between 1970-1979, *Ralstonia* remained dominant at 45%, with *Rothia* (7.9%) and *Staphylococcus* (7.8%). In libraries prepared from specimens collected between 1980-1989, assemblages were dominated by *Terasakiella* (43%), *Neptuniibacter* (13%), and *Vibrio* (4.8%). Finally, in libraries prepared from specimens collected between 2010-2019, 16S rRNA amplicons were more diverse, with *Ralstonia* (11%), *Vibrio* (10%), and *Photobacterium* (6.3%) among the most prevalent taxa.

The facility, year and decade of collection, as well as DaScPc18S rRNA detection have significant correlations with genera representation in our samples (Figure 4). The facility in which the samples were stored has a significant positive correlation (Pearson Correlation, R^2^ = 1, p-val < 0.05) with *Fusobacterium*, *Neisseria*, and *Roseomonas* genera, as well as a significant negative correlation (Pearson Correlation, R^2^ = −1, p-val < 0.05) with *Veillonella*, and *Pseudonocardia*. The year when the sample was collected also had significant positive correlations (Pearson Correlation, R^2^ = 1, p-val < 0.05) with *Veillonella*, and significant negative correlation (Pearson Correlation, R^2^ = −1, p-val < 0.05) *Fusobacterium*, *Neisseria*, and *Roseomonas* genera. When agglomerated into decade of collection, with the goal of identifying major changes across time, the same groups as for the year of collection were significant, with the addition of a significant negative correlation with *Limibacillus*. Presence of DaScPc 18S rRNA, also yielded significant positive correlations (Pearson Correlation, R^2^ = 1, p-val < 0.05) with *Fusobacterium*, and *Neisseria;* and significant negative correlation (Pearson Correlation, R^2^ = −1, p-val < 0.05) with *Veillonella*, *Pseudocardia*, HOC36 (Gammaproteobacterium), and *Kiritimatiellota.* Despite these correlations, the facility specific difference between samples prior to 1980 and after 1990 impair our ability to assess what is a biological shift or sample management and storage bias.

**Figure 4.**
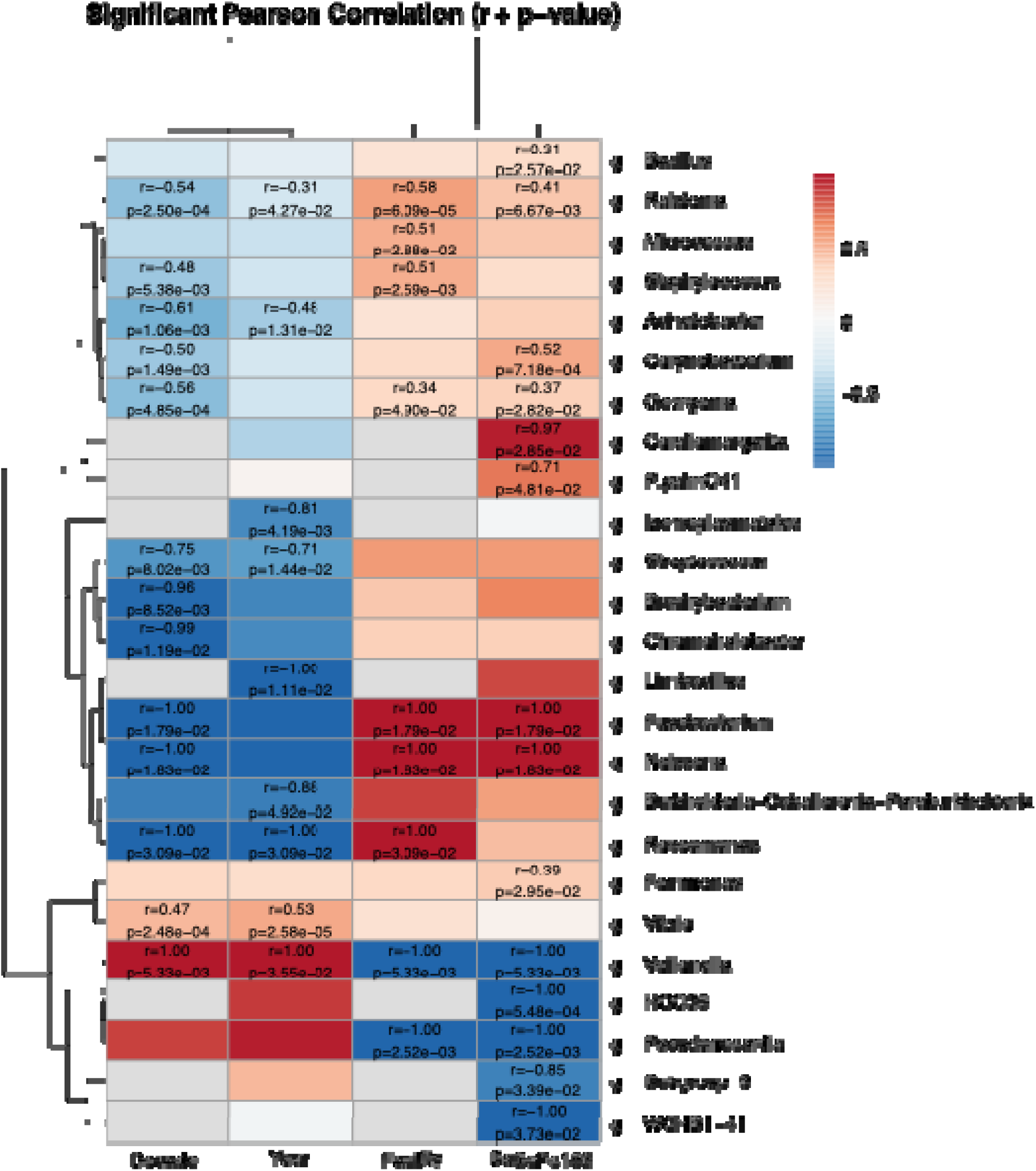
Heatmap shows correlations, based on Pearson Correlation Coefficient, between genera and environmental factors.

## Discussion

Our results suggest that we can detect Philaster-like ciliates across a wide temporal range using 18S rRNA, yet the usage of 28S rRNA showed, species or more accurate genus level detection, was less successful. Museum specimens serve as genomic archives for tracking the presence of environmental pathogens, offering unique insights into past disease dynamics (Raxworthy & Smith, 2021). The 18S rRNA gene is widely used for phylogenetic studies in eukaryotes to identify distinct species, yet studies exploring ciliate genetic markers have shown that morphologically distinct ciliates can share identical 18S rRNA genes, and that the ITS and 28S rRNA provide a better resolution for species identification and phylogenetic divergence (Fan et al., 2021; Somasundaram & Yu, 2025). Despite the benefits of using the 28S rRNA for lower-level taxonomic classification (e.g. species and genus), its ability to acquire insertions and deletions in expansion segments, can also render it too distinct when comparing samples across large time scales, potentially limiting our ability to identify the ciliate with primers derived from contemporary DaScPc 28S rRNA sequences (Pereira & Baldwin, 2016; Somasundaram & Yu, 2025). Our combined approach, with varying degrees of success, was able to recover DaScPc 18S rRNA from a variety of specimens over time, but only related (not identical) 28S rRNA sequences from two *D. antillarum* specimens in 1982 (earlier than mass mortality) and 1983 (during mass mortality), both from grossly normal specimens at then-unaffected sites (Lessios, Robertson & Cubit, 1984). None of the tested museum specimens amplified when using the 5.8S/ITS region. Furthermore, detection of DaScPc in only single urchins amongst multiple from the same collection (i.e. multiple specimens per jar) suggests that these are heterogeneously present in urchin microbiomes, but we cannot discount the possibility of cross-contamination between urchins within jars. The lack of identical 28S rRNAs to DaScPc in two specimens and lack of amplification altogether across all other DaScPc 18S rRNA-positive specimens suggests that 18S rRNA alone may be an insufficient marker for DaScPc in environmental or echinoid samples. The DaScPc 18S rRNA detections in museum specimens may represent a related Philaster ciliate that given the grossly normal state of the specimens, were most likely not participating as a pathogenic agent. These results aide in our understanding the ecology of DaScPc-like ciliates and inform future work targeting detection of DaScPc as it is associated with disease, vectors, and environmental reservoirs. Additionally, it underscores the limitations posed by DNA fragmentation due to preservation or that which can be induced due to column-based DNA extraction protocols which can hinder sequence amplification success (Höpke et al., 2019; Velasco-Cuervo et al., 2019). More work is required to improve on genomic material recovery and preservation storage influence on contamination and its effects of sequencing efforts.

Microbial community analysis revealed there was no representation of target bacterial biomarkers known to be associated with DaSc infection. The microbiome composition of preserved *D. antillarum* specimens may provide insights into both historical microbial diversity and the effects of long-term preservation on microbial DNA. Previous work on non-preserved specimens confirms our findings showing no significant dissimilarities across *D. antillarum*’s epibiota throughout different geographical expanses (Rodríguez-Barreras et al., 2023). Our data revealed distinct facility-specific microbial signatures and notable temporal shifts in bacterial communities, suggesting that collection conditions and age in the collection may influence the microbiome. This contradicts previous studies that saw no correlation between collection time and the microbial community composition (Chalifour, Elder & Li, 2022). Recent studies exploring the microbiome of *D. antillarum*, on non-preserved specimens, show similar dominant genera than those seen in the 2010-2019 preserved specimens (Rodríguez-Barreras, Tosado-Rodríguez & Godoy-Vitorino, 2021; Rodríguez-Barreras et al., 2023; Vilanova-Cuevas et al., 2025a). For example, high relative abundance of *Propionigenium* in more recent decades (2010-2019) aligns with recent studies that establish this genus as part of the microbiota of grossly normal *D. antillarum* in non-preserved specimens (Rodríguez-Barreras, Tosado-Rodríguez & Godoy-Vitorino, 2021; Rodríguez-Barreras et al., 2023; Vilanova-Cuevas et al., 2025a). Several genera that varied across collection periods, such as *Corynebacterium*, *Bacillus*, *Staphylococcus*, and *Rothia*, are spore-forming Gram-positive bacteria, which may be more resistant to long-term preservation decay due to their greater structural integrity and resistance to ethanol-induced lysis and DNA fragmentation than Gram-negative bacteria (Duquenoy et al., 2020). This could contribute to their persistence in older specimens and may partially explain biases in microbial profiles recovered from preserved samples. Such preservation-related effects complicate efforts to distinguish true ecological shifts from the significant correlations seen in our study based on facility and collection period.

Despite the possible bias, our analysis revealed high abundance of several Gram-negative genera, including *Terasakiella*, *Neptuniibacter*, and *Vibrio*, in samples collected between 1980-1989, which were also highly abundant in DaScPc-infected *D. antillarum* from the 2022 mass mortality (Vilanova-Cuevas et al., 2025a). These genera are commonly associated with marine environments and invertebrate hosts, and their detection across preserved and fresh samples suggests ecological relevance and potential stability over time (Rodríguez-Barreras et al., 2023; Shaw et al., 2023; Vilanova-Cuevas et al., 2025a). Moreover, our analysis showed significant difference between presence and absence of DaScPc 18S rRNA sequences, but none of the identified bacterial amplicon sequences match the Diadematidae scuticociliatosis biomarker, *Fangia hongkongensis*, from the 2022-2023 mass mortalities (Vilanova-Cuevas et al., 2025a). These findings highlight the complexity of interpreting microbial shifts in preserved collections, where both ecological and preservation-related factors may shape microbial community profiles.

## Conclusion

The causative agent of *D. antillarum* mass mortality from 1983-1984 remains unknown, as will likely be the case into the future, due to the lack of affected specimens for interrogation with modern molecular approaches. Nonetheless, studying museum preserved specimens of grossly normal *D. antillarum* in this study identified 18S and 28S rRNA sequences of DaScPc or a closely related ciliate for over 50 years. This finding suggests Philaster ciliates potentially related to DaScPc may be normally associated with *D. antillarum*, since these were recovered from grossly normal specimens. This builds upon a wider understanding that this clade also occurs on sympatric corals and other surfaces where urchins typically reside. Our study showcases the utility of museum specimens for investigating presence or absence of current pathogenic agents, identification of bacterial biomarkers, and studying microbial communities over time, while also highlighting the difficulties of possible biotic and abiotic biases at play. Future work focused on microbial community dynamics in museum specimens should test a wider range of the collection to identify possible contaminants associated with the facility and storage conditions and address those biases bioinformatically to retain biological information relevant to the sample.

## Acknowledgements

The US National Museum of Natural History (Smithsonian), Institution Department of Invertebrate Zoology and the Florida Marine and Wildlife Conservation Commission thanked for providing samples used in this research.

---

**Supplementary Table 1.**
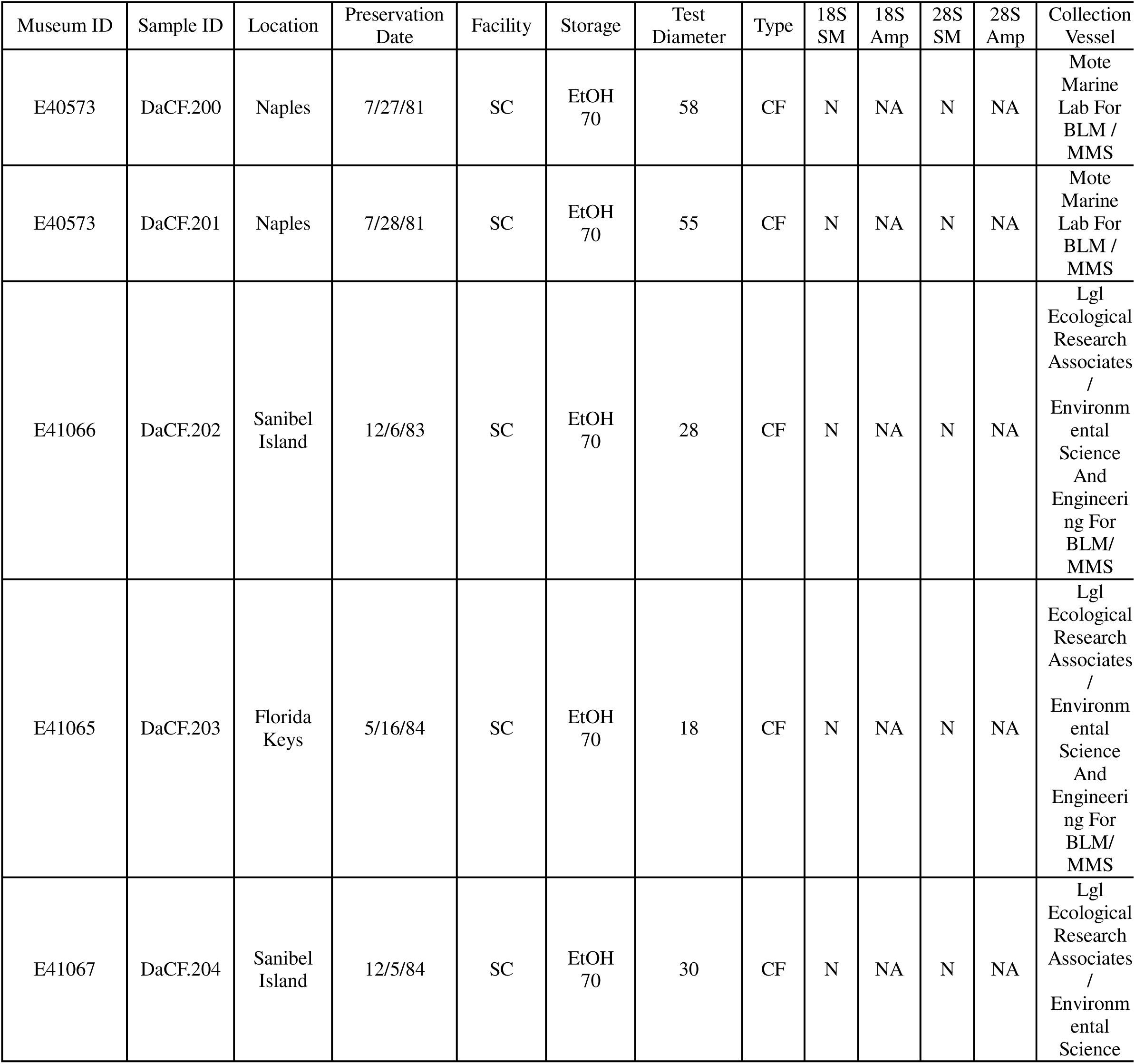
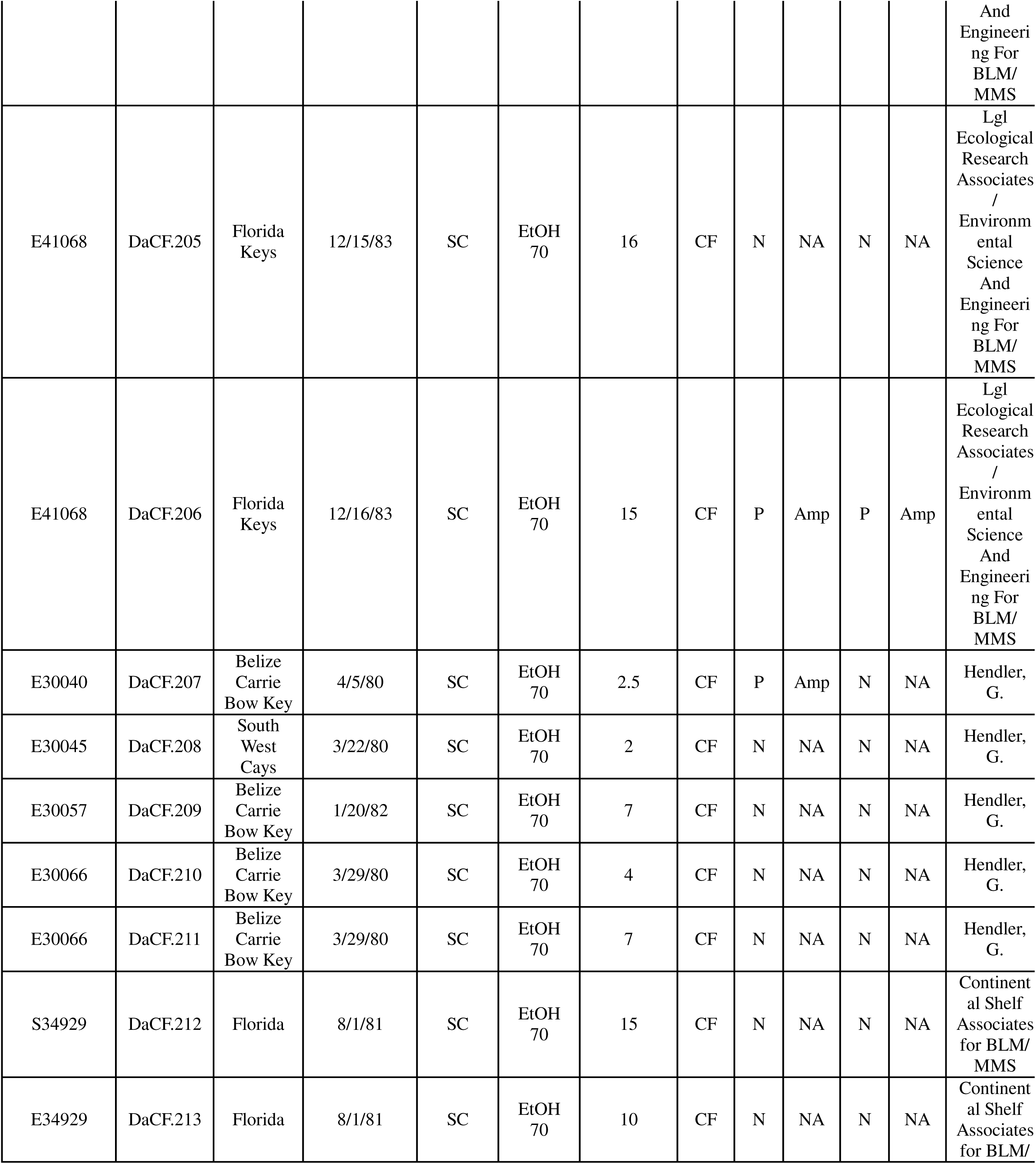
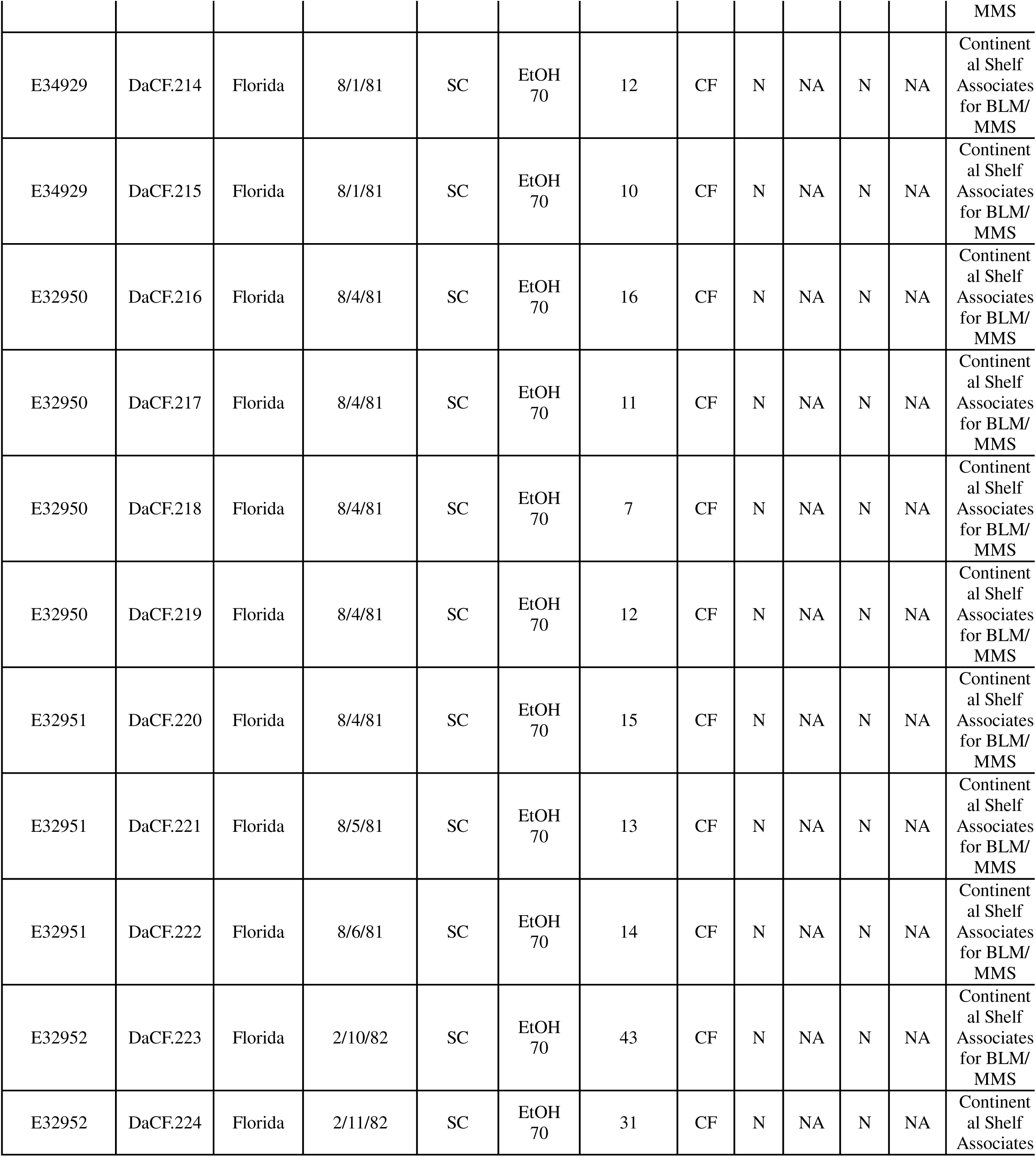
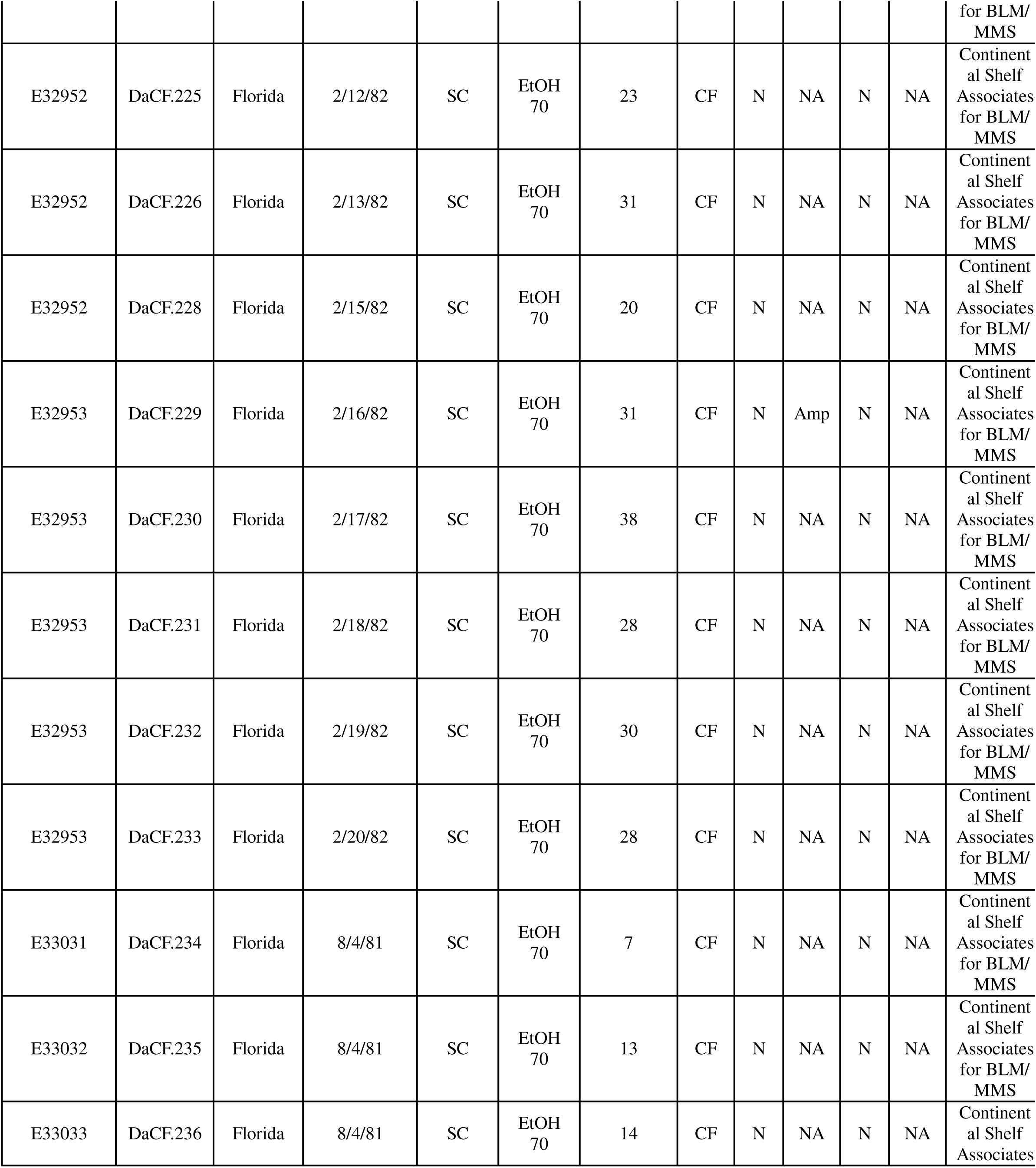
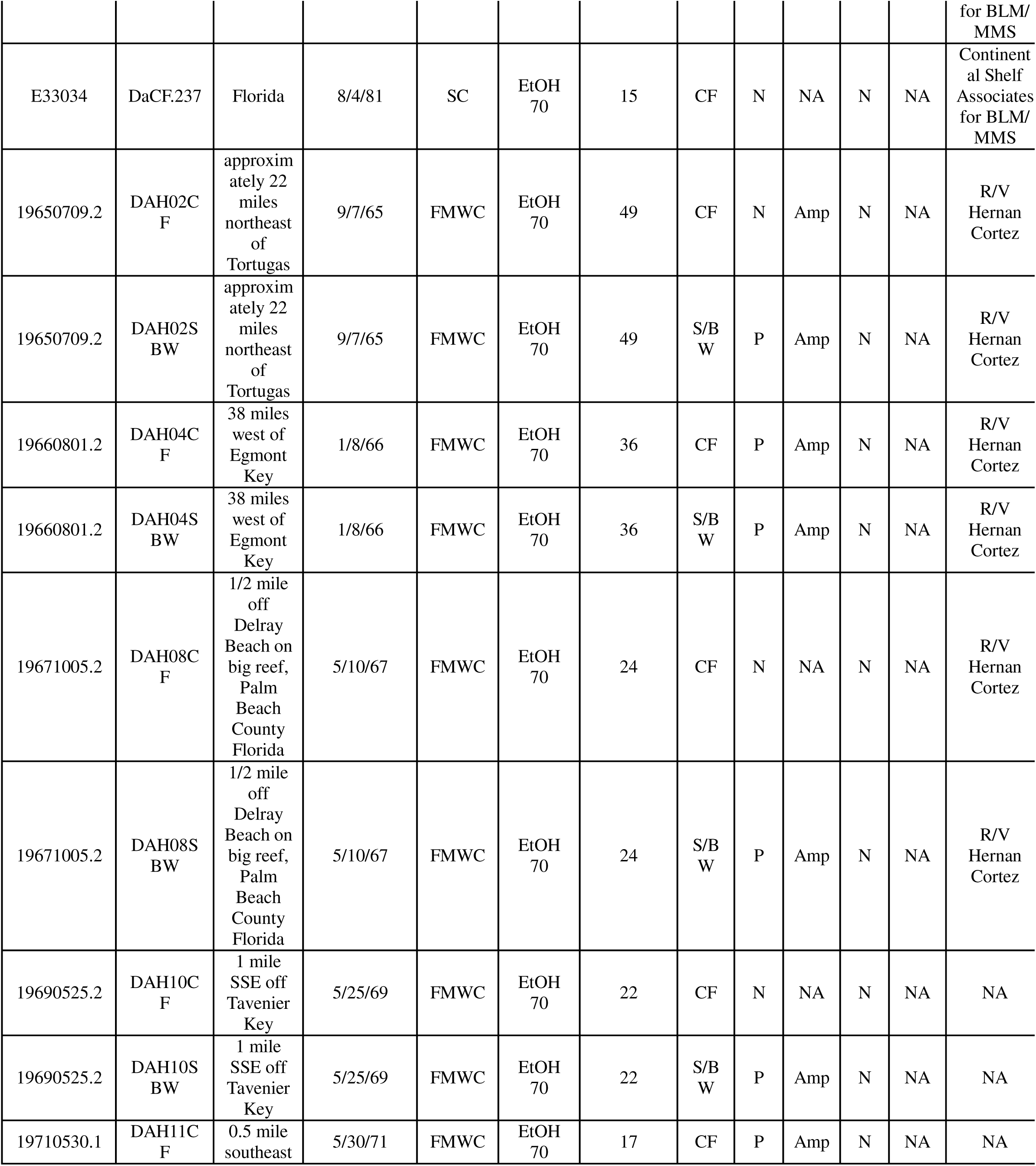
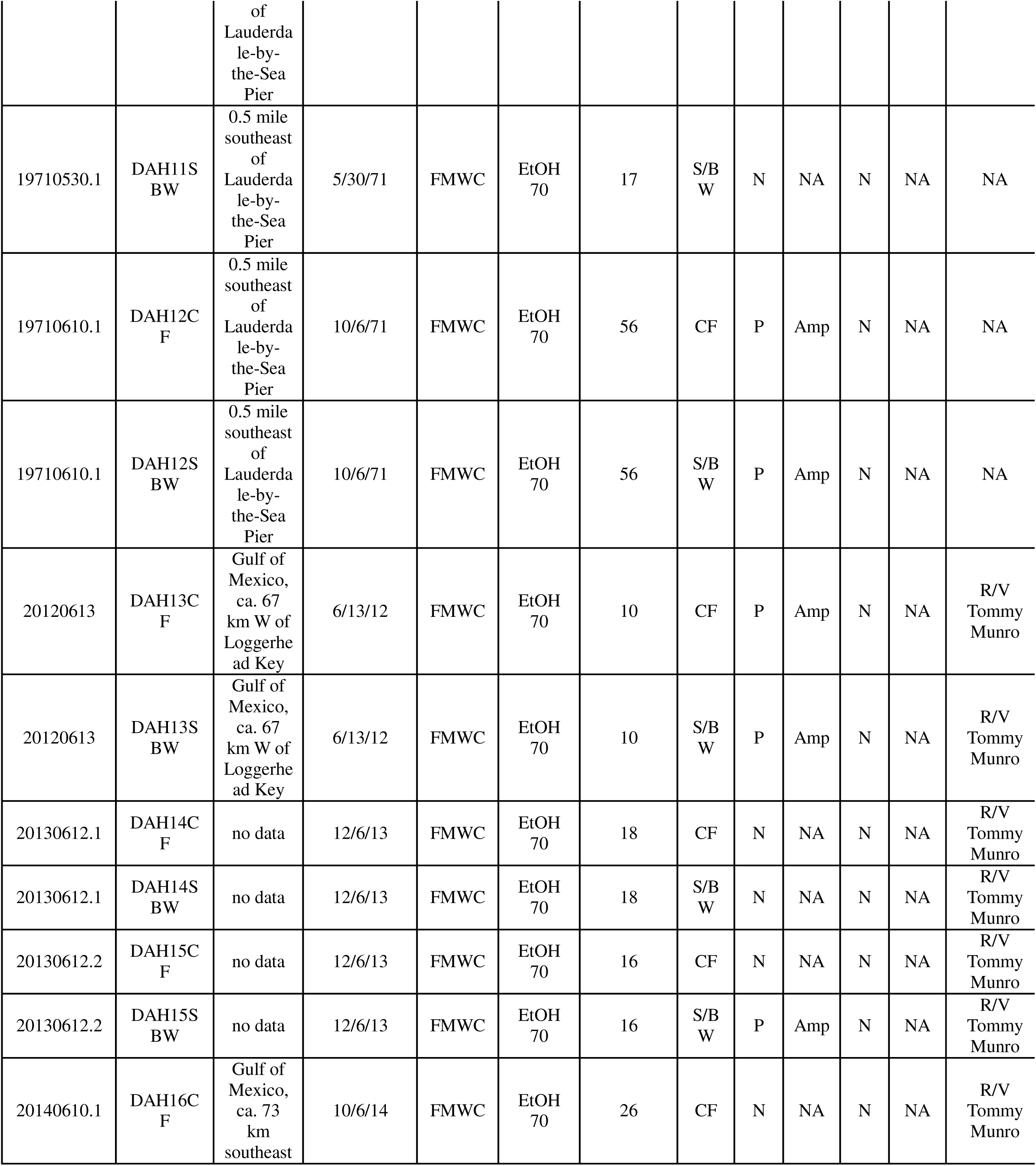
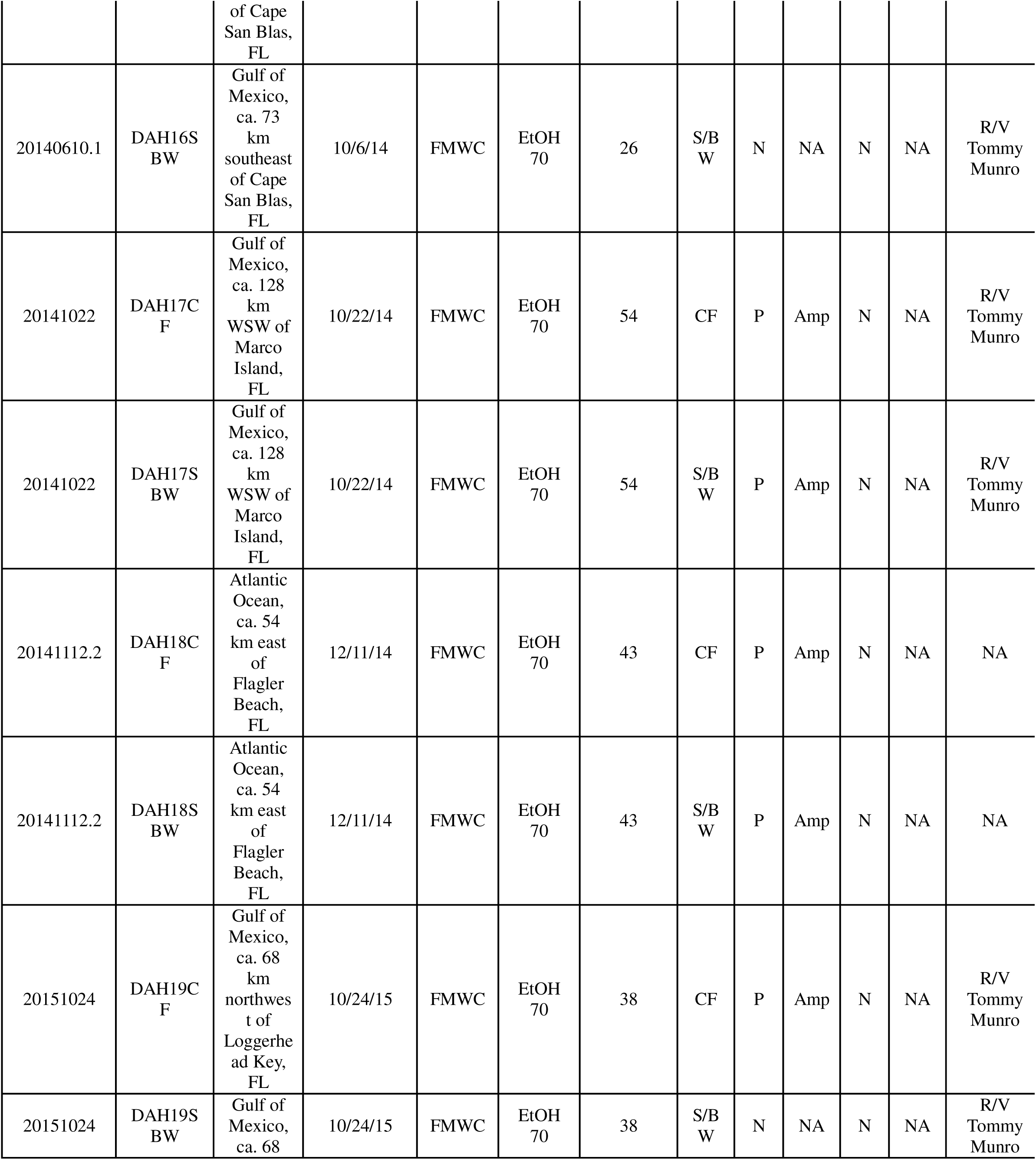
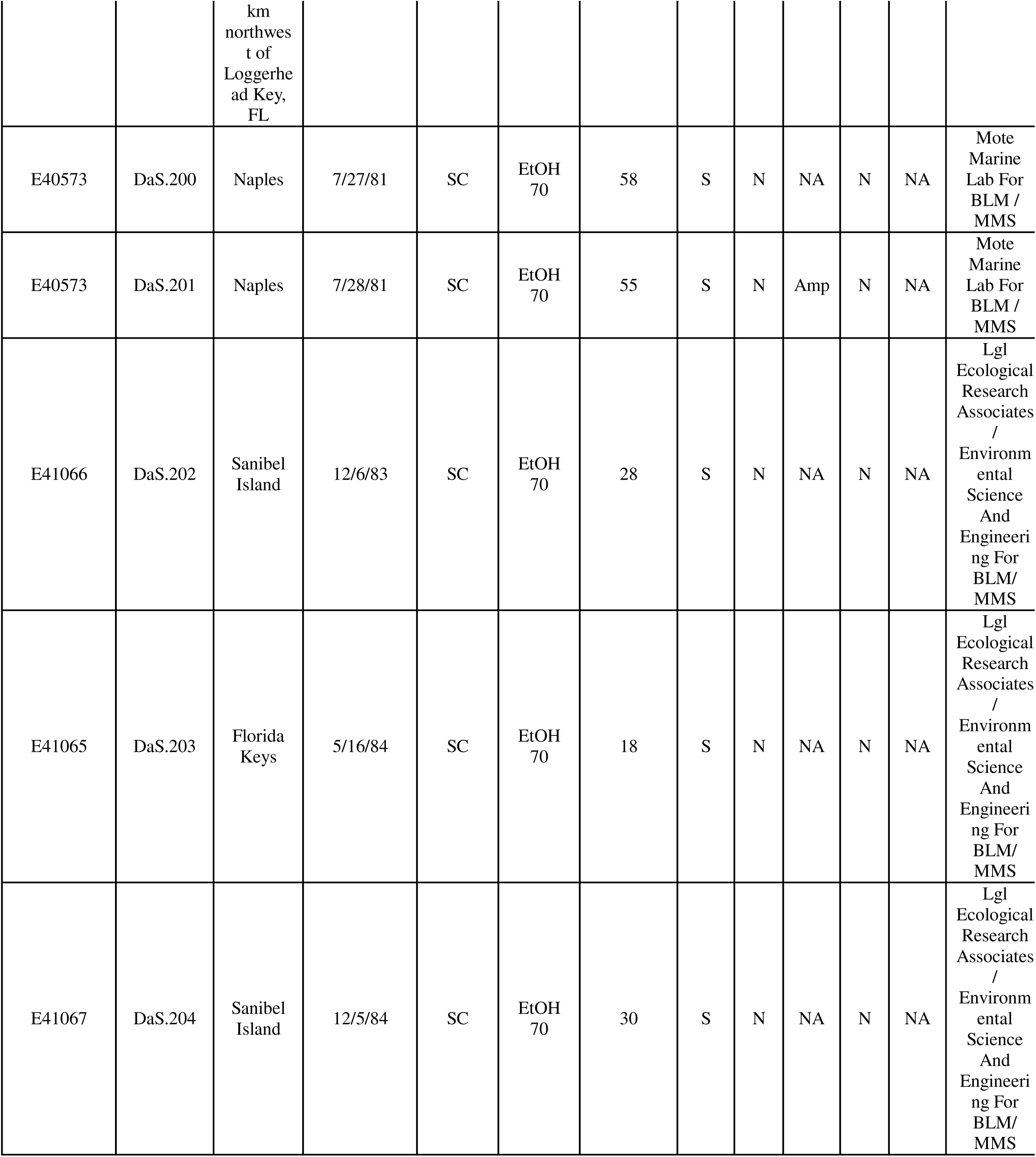
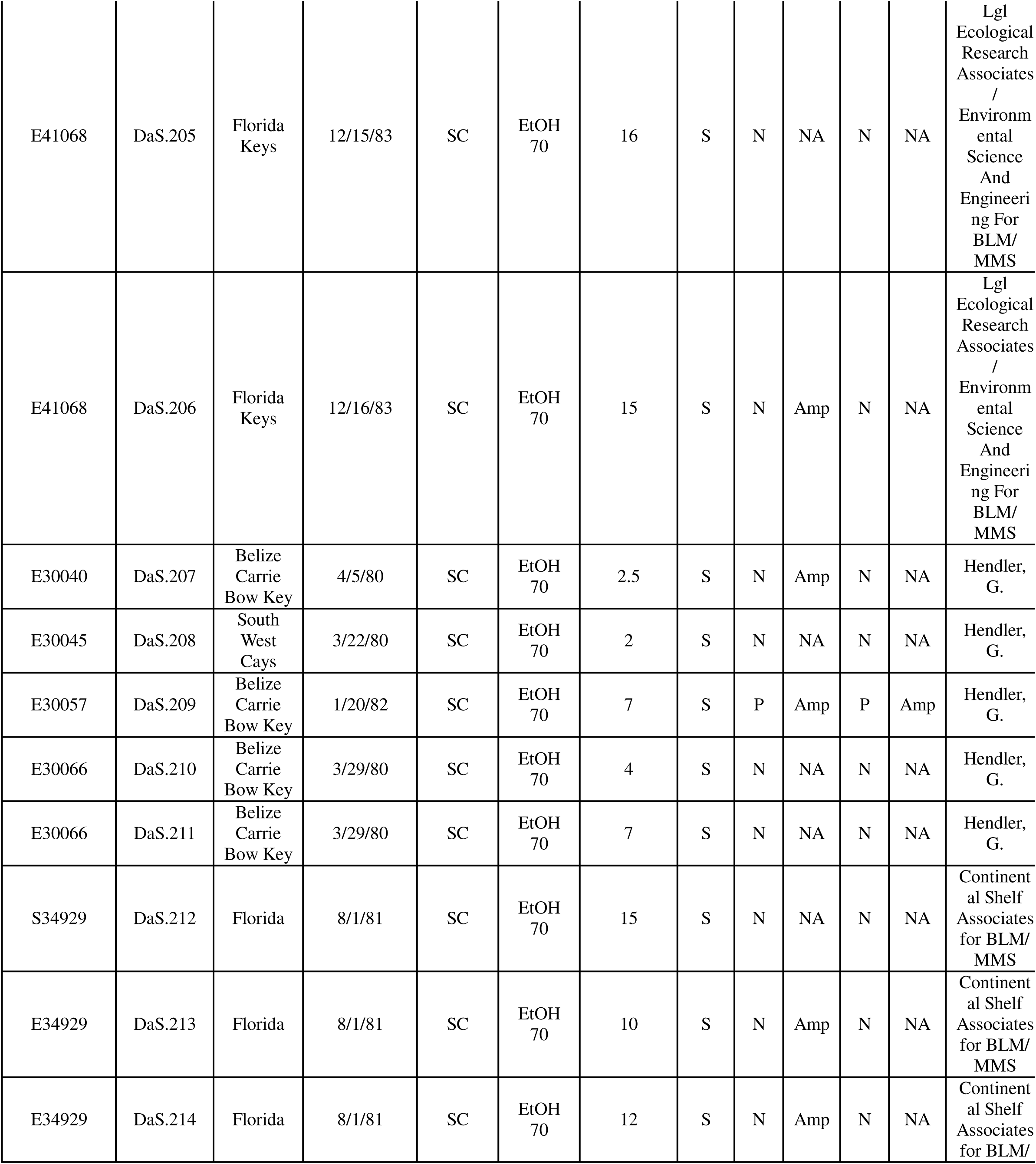
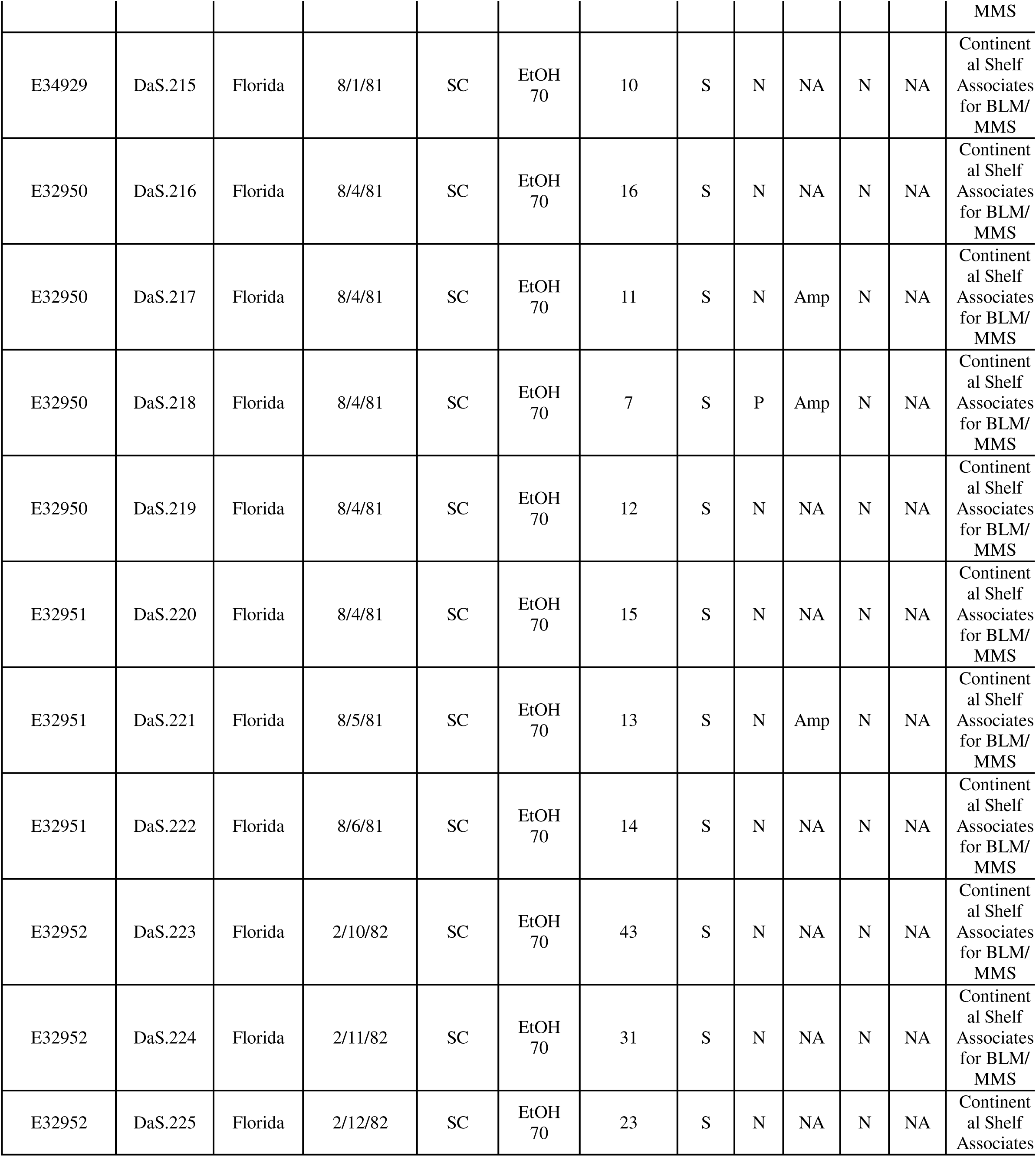
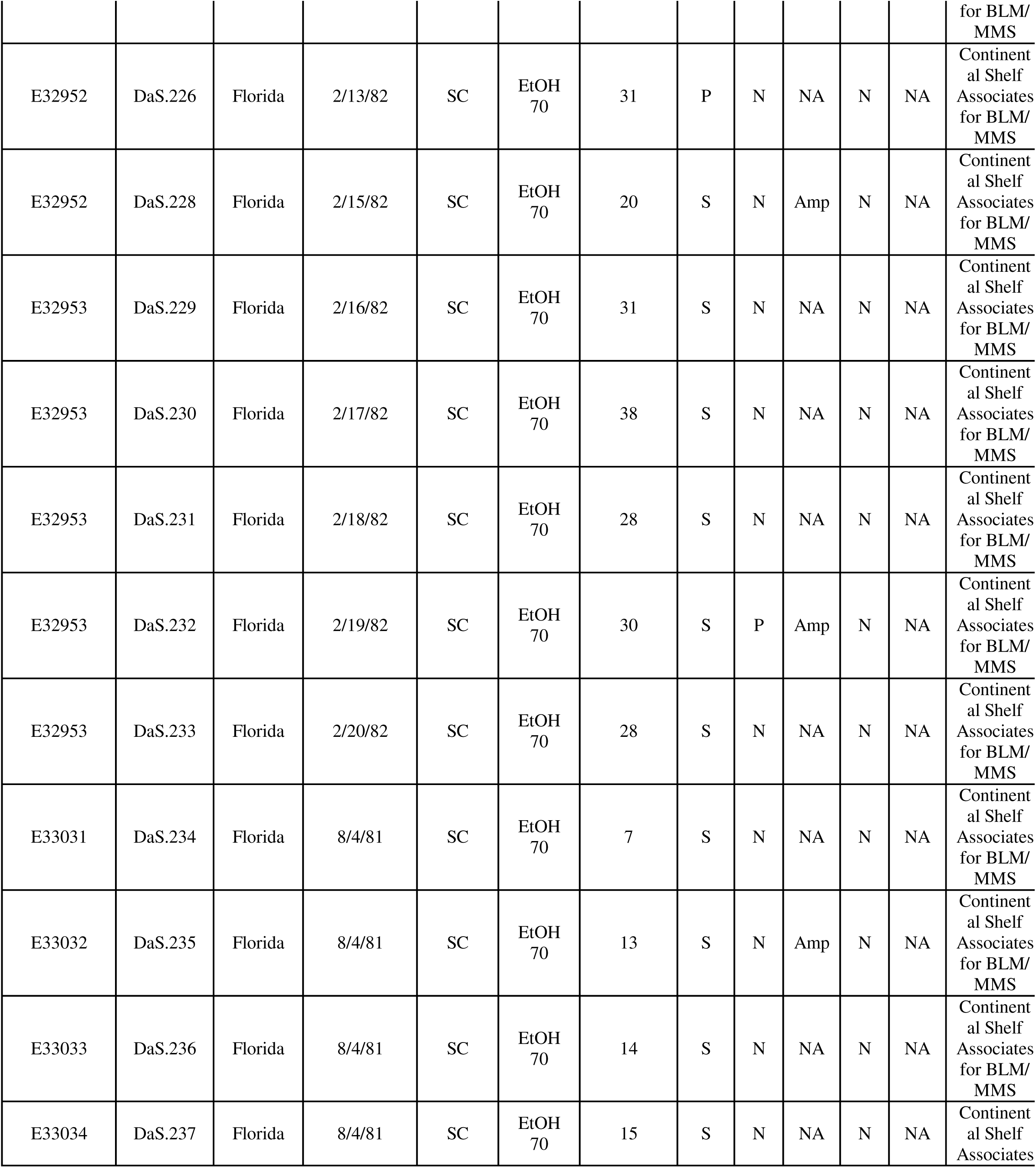
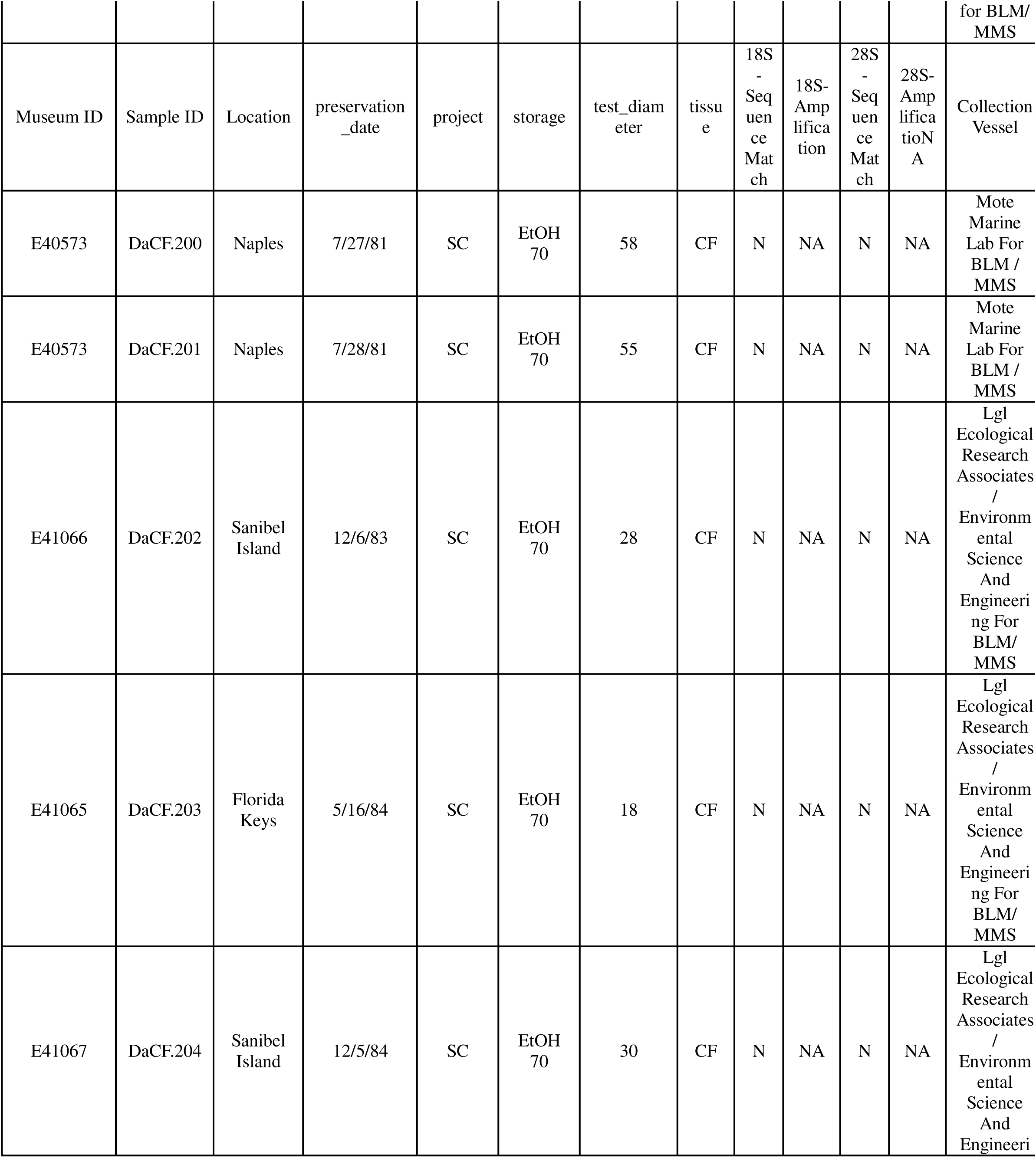
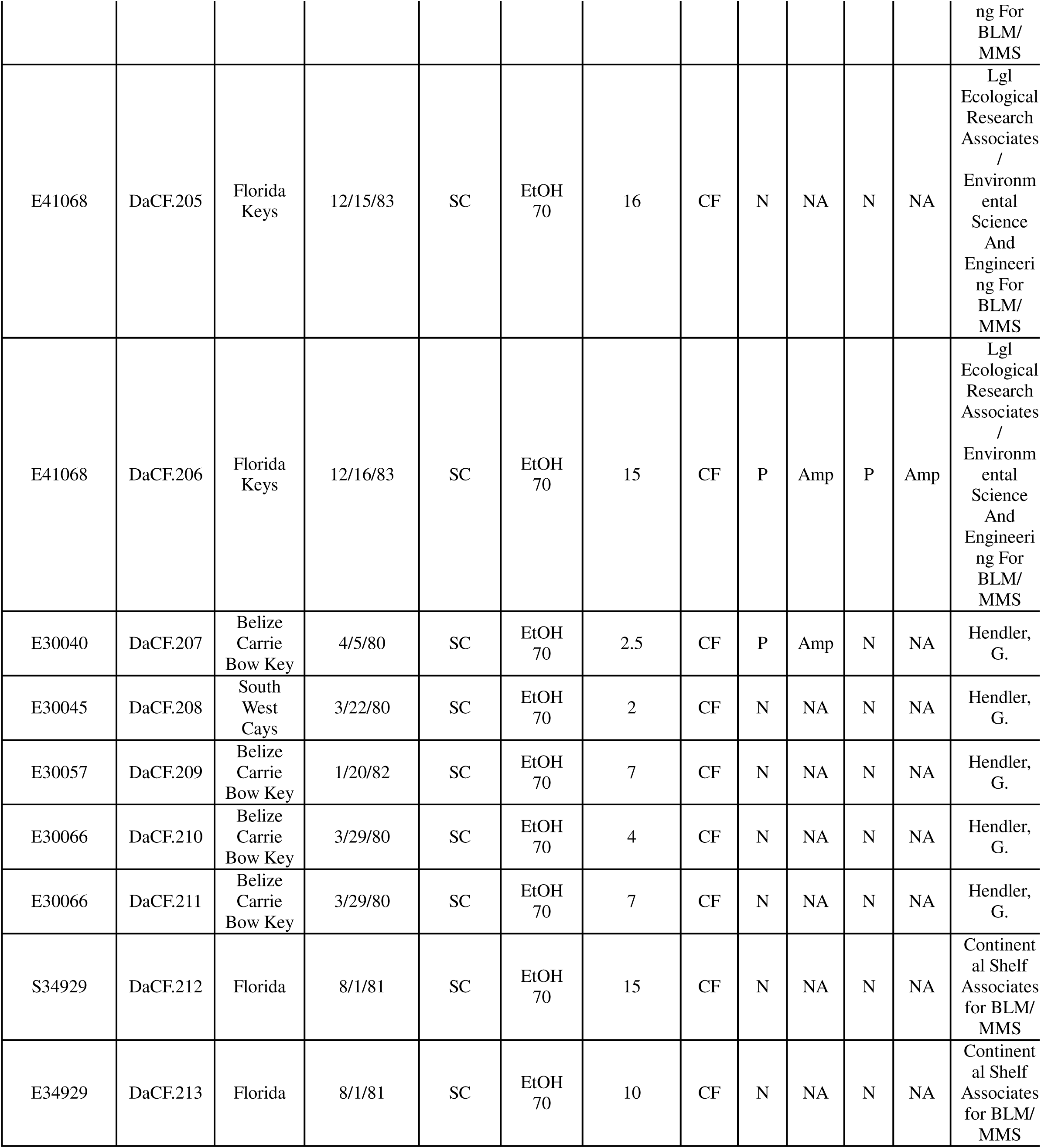
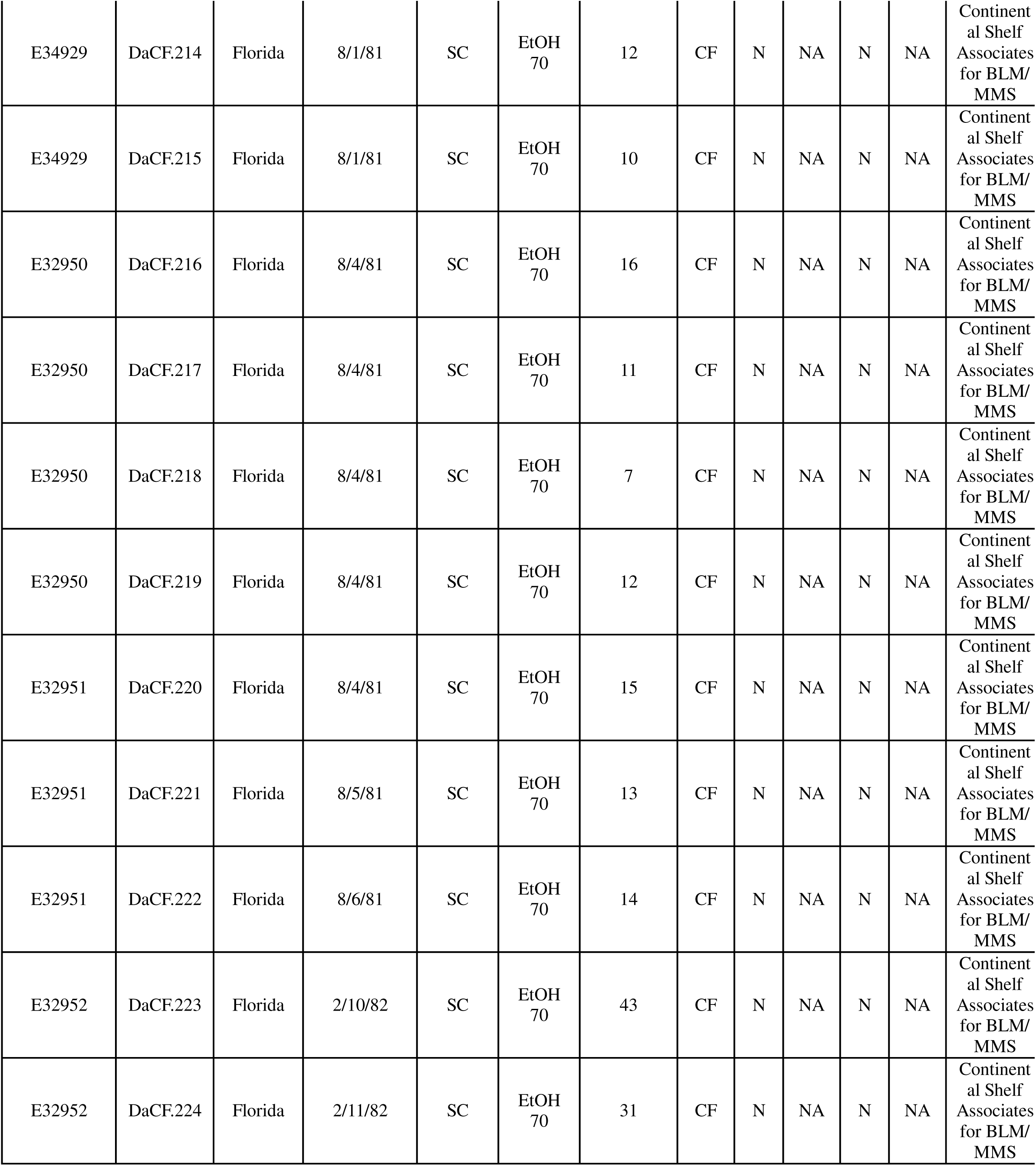
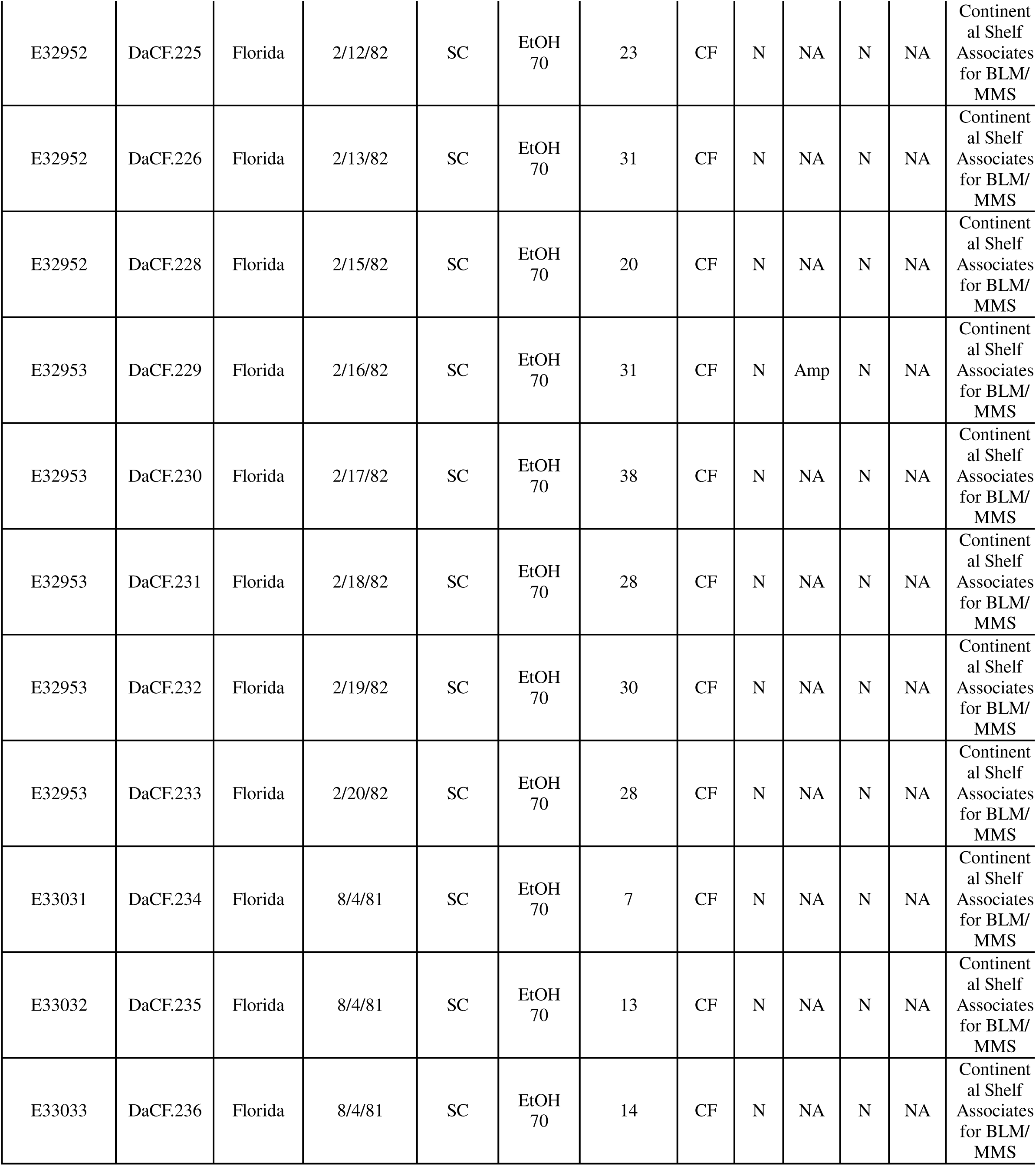
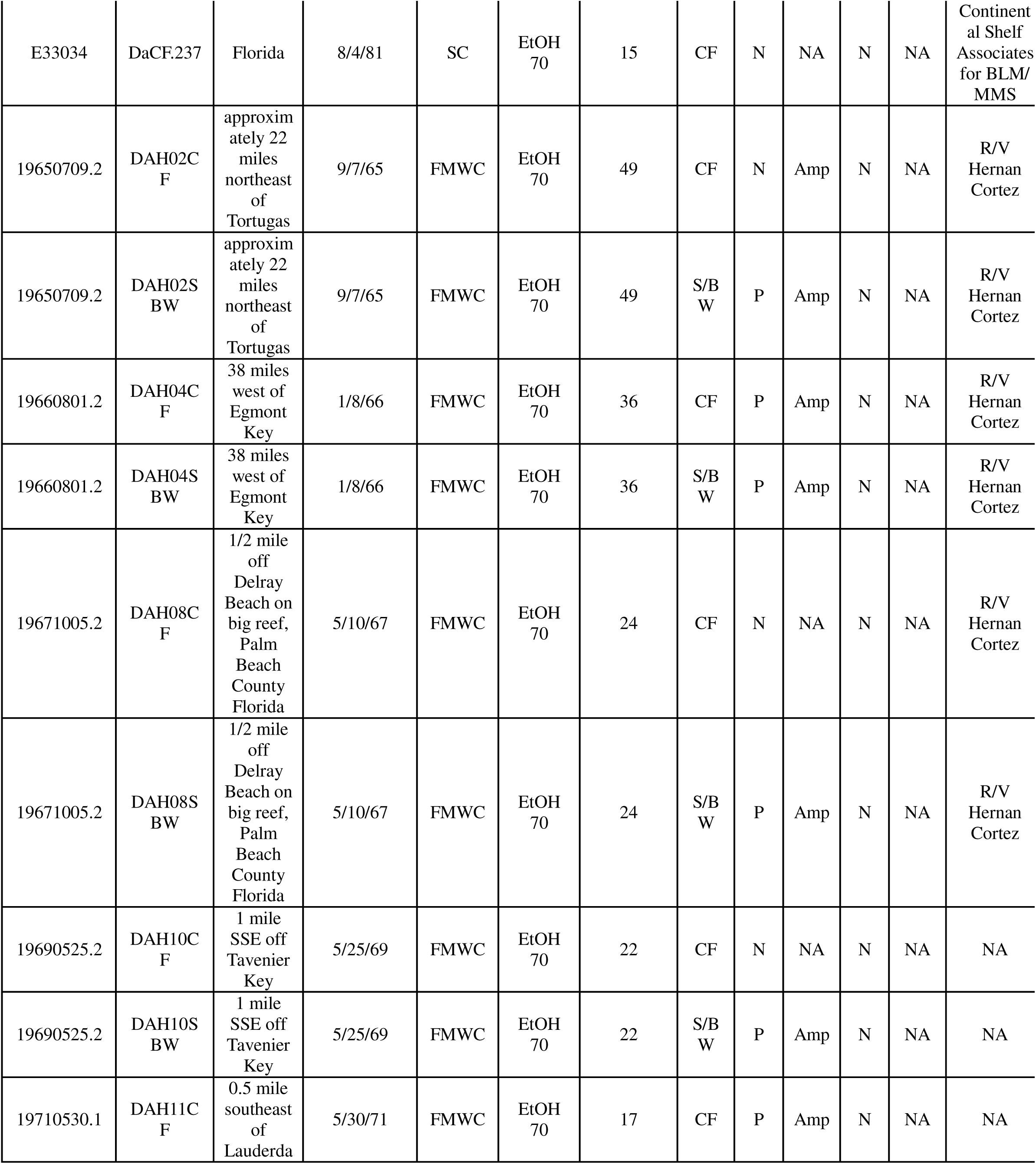
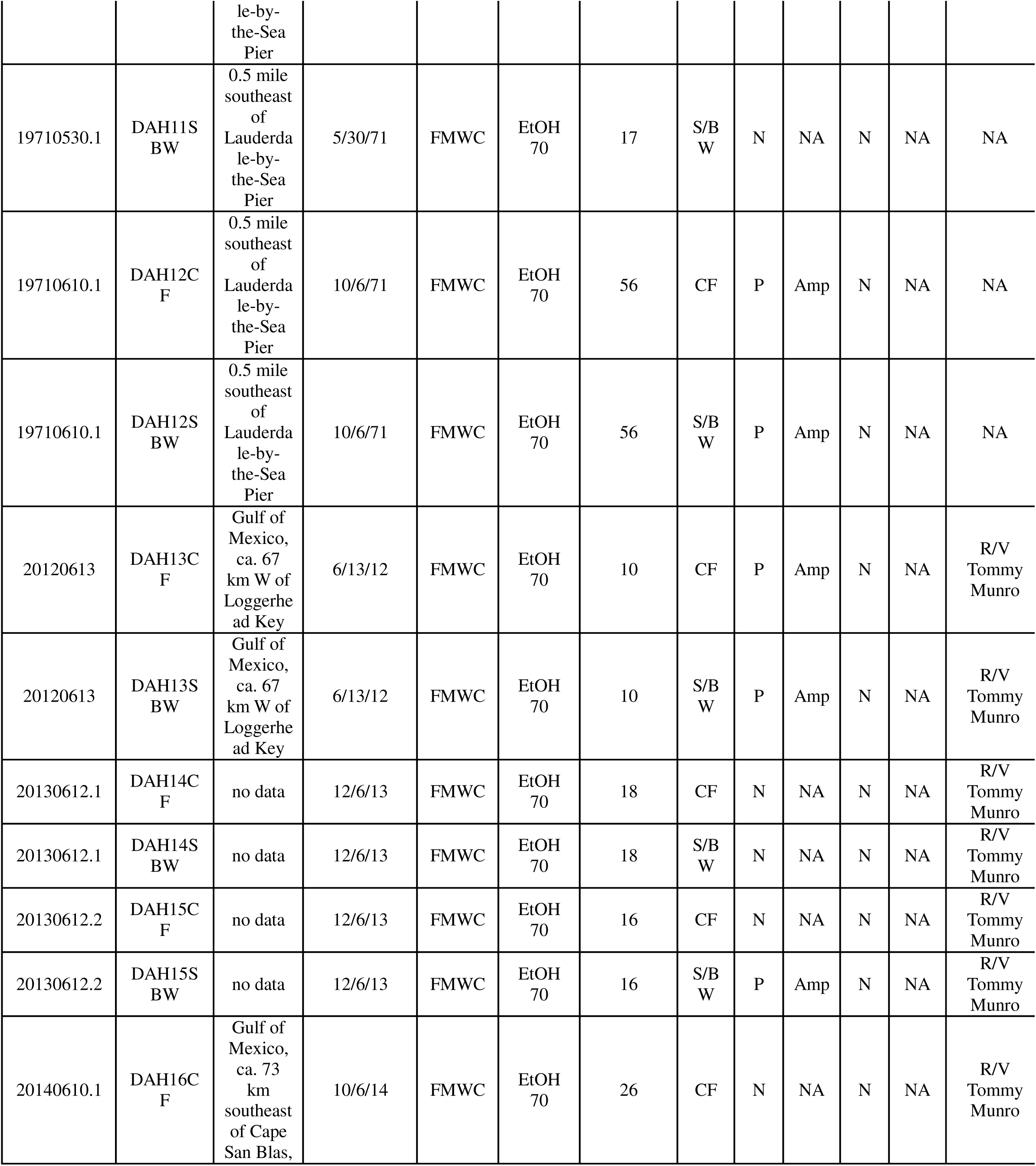
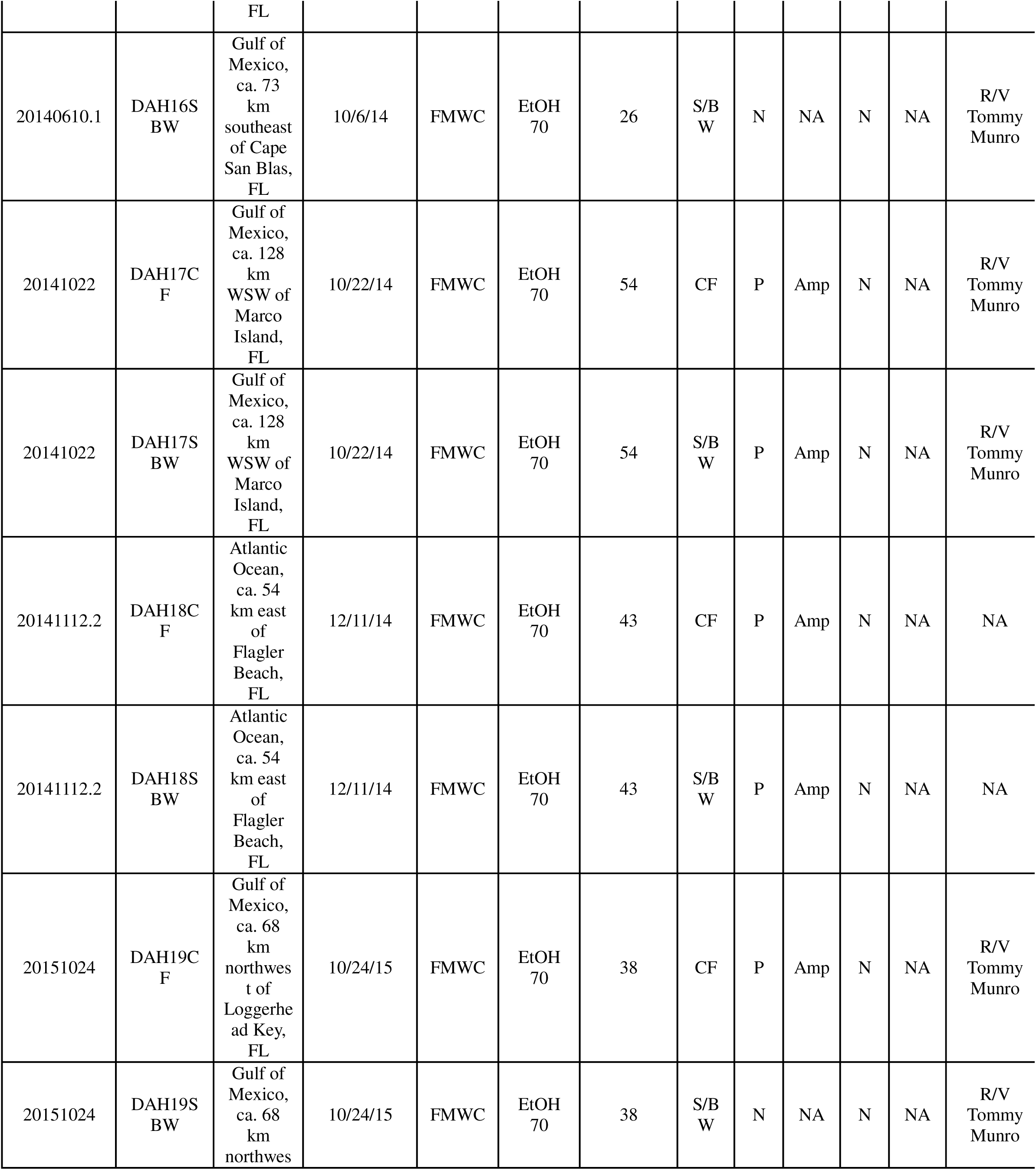
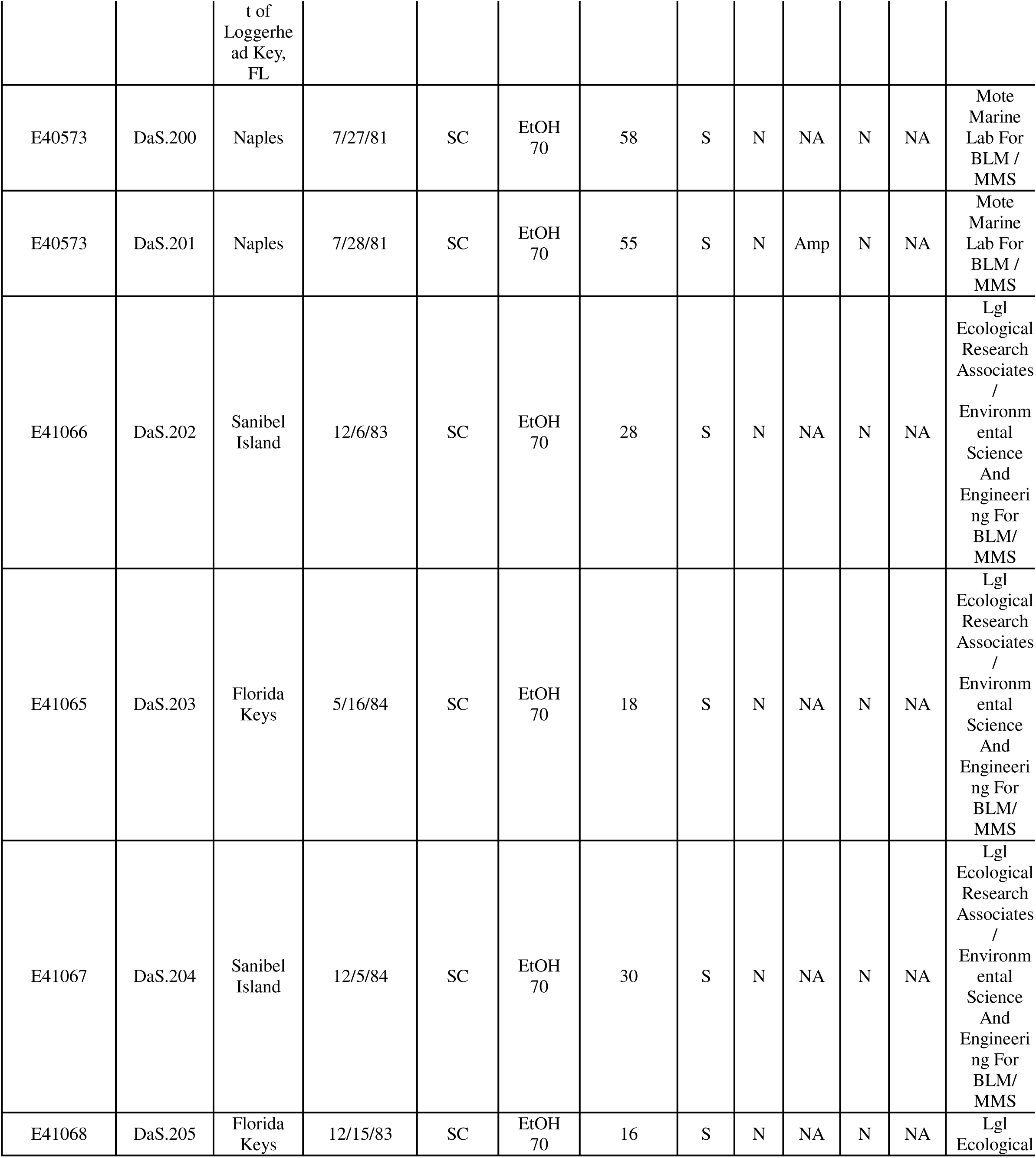
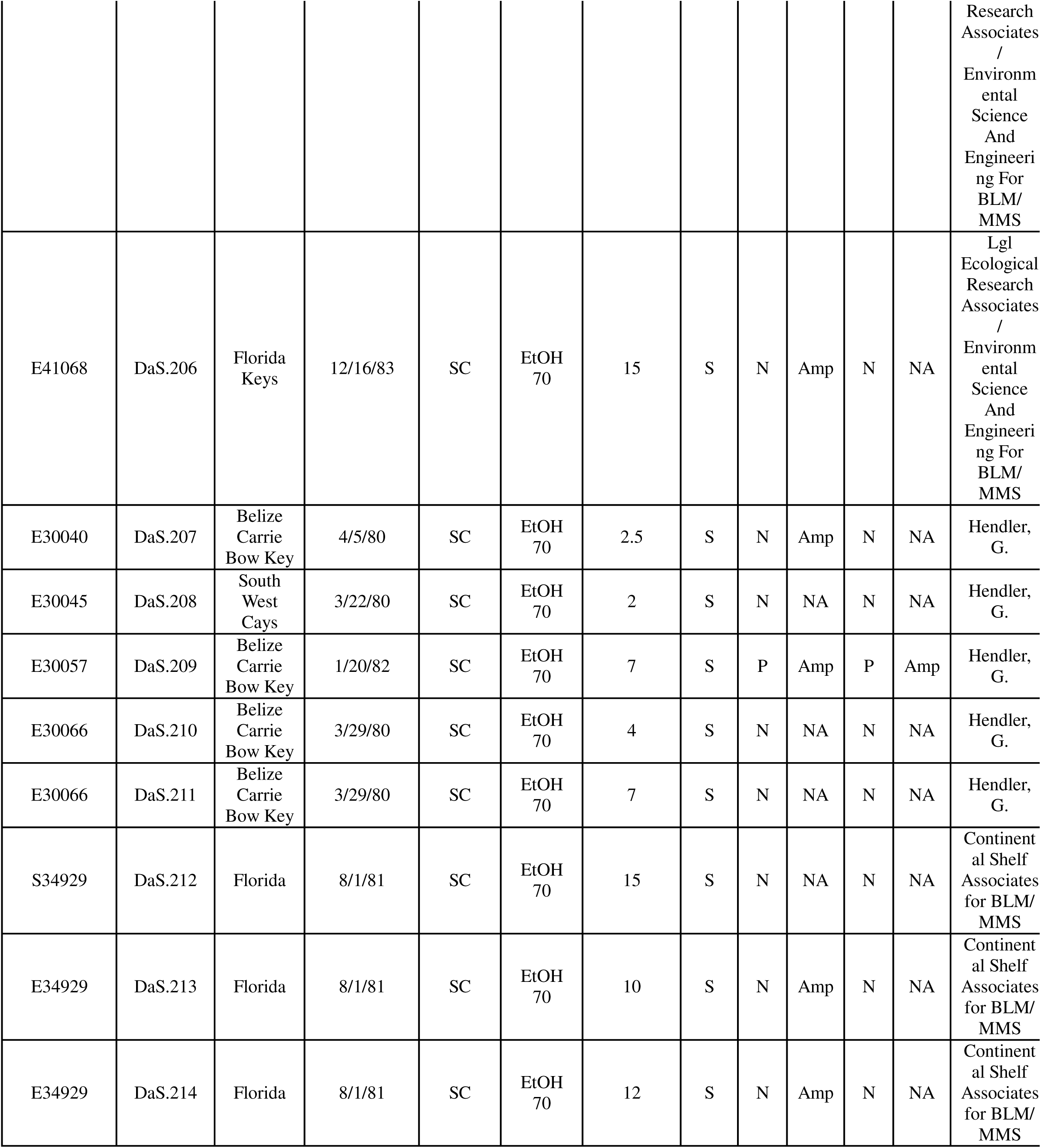
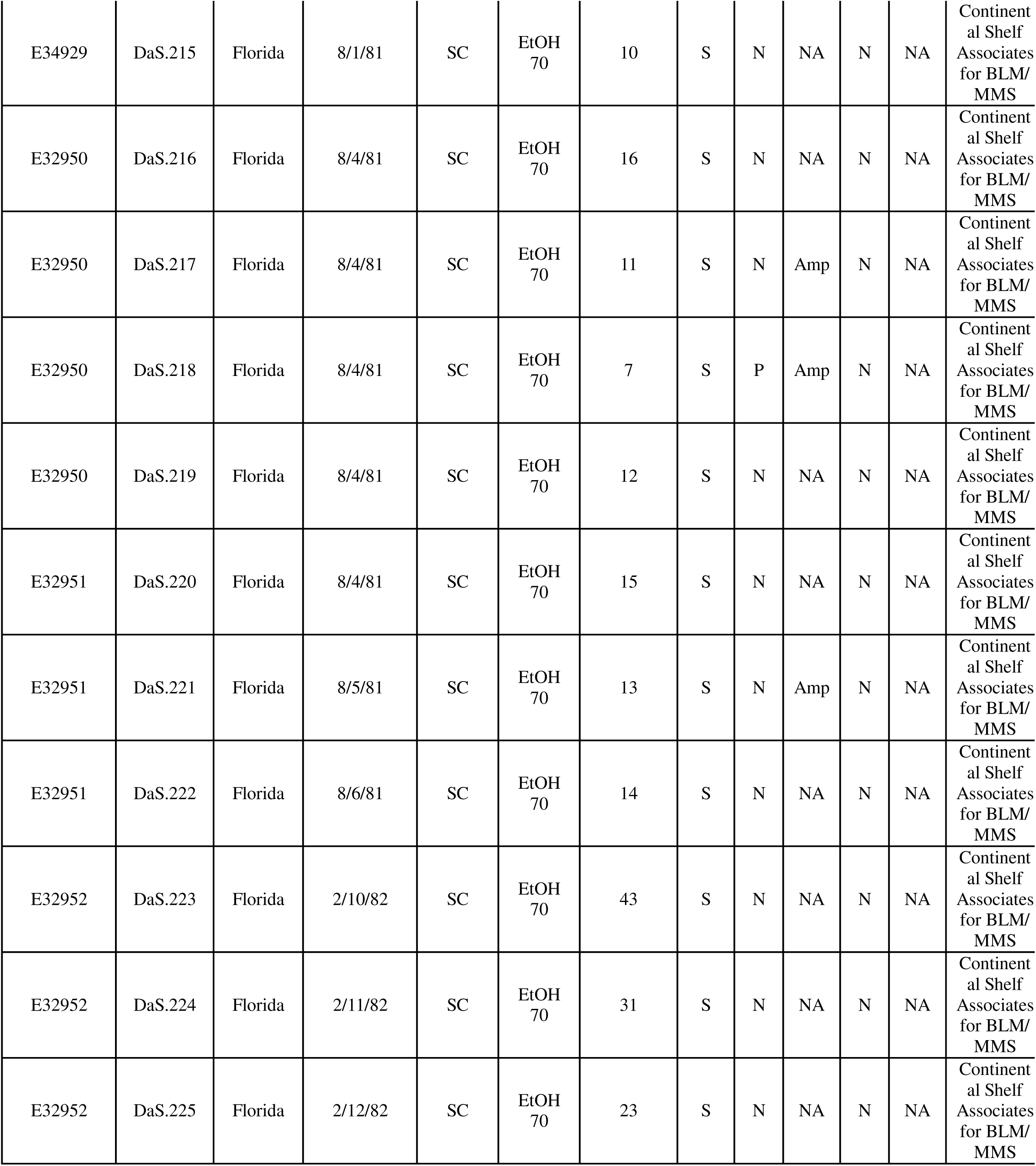
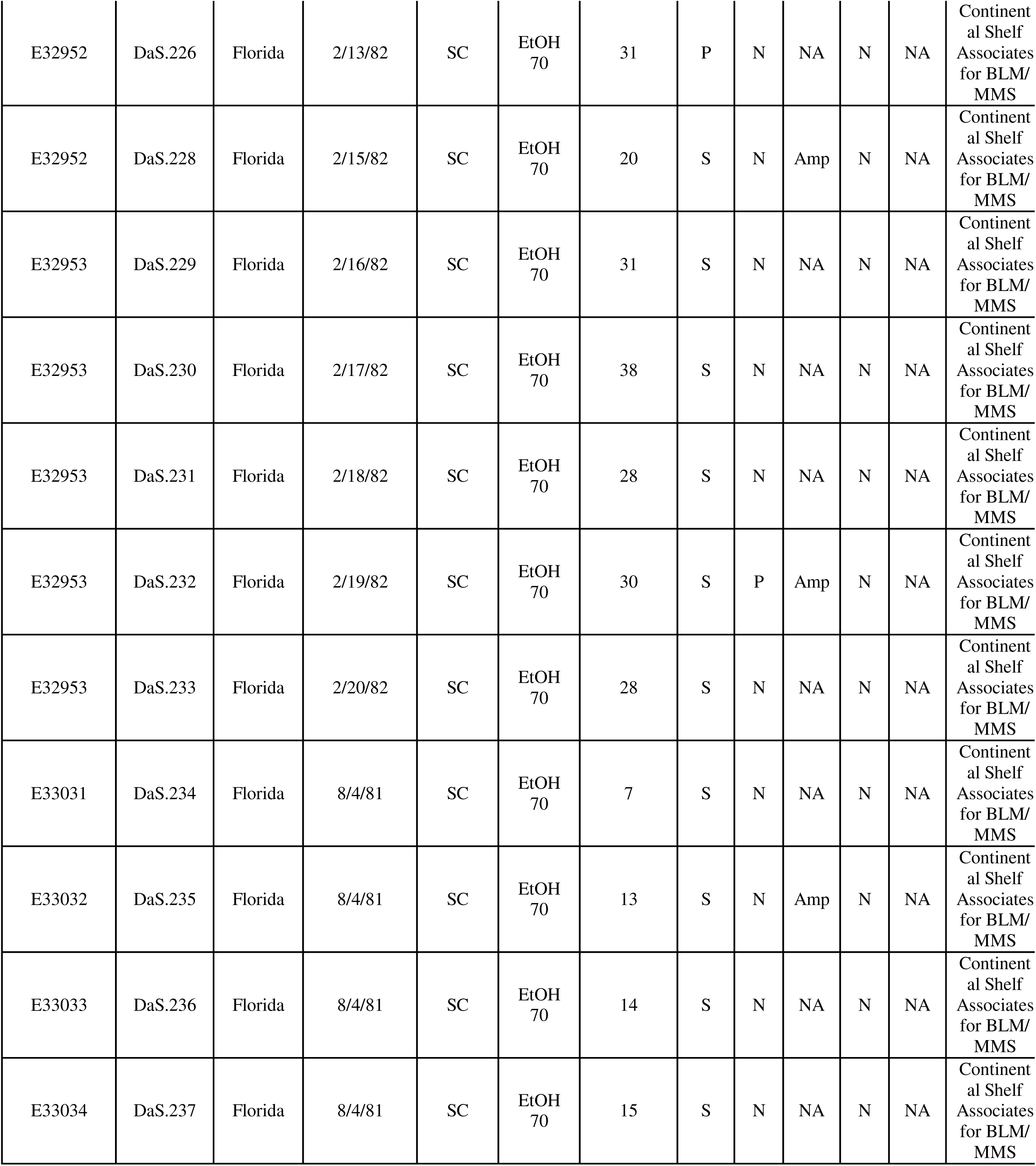
Metadata associated with all the analyzed samples is below, museum ID refers to the code name given to an individual sample by the host facility. Facilities have been abbreviated to SC for Smithsonian Collection and FMWC for Florida Marine and Wildlife Commission. The Sample Type has been abbreviated CF – coelomic fluid, S – spine, and S/BW – spine and body wall. Column headers have also been abbreviated for 18S and 28S sequence match (SM), and amplification (Amp) where N – Negative, NA – No amplification, Amp – amplified, and P – positive.

**Supplementary Figure 1.**
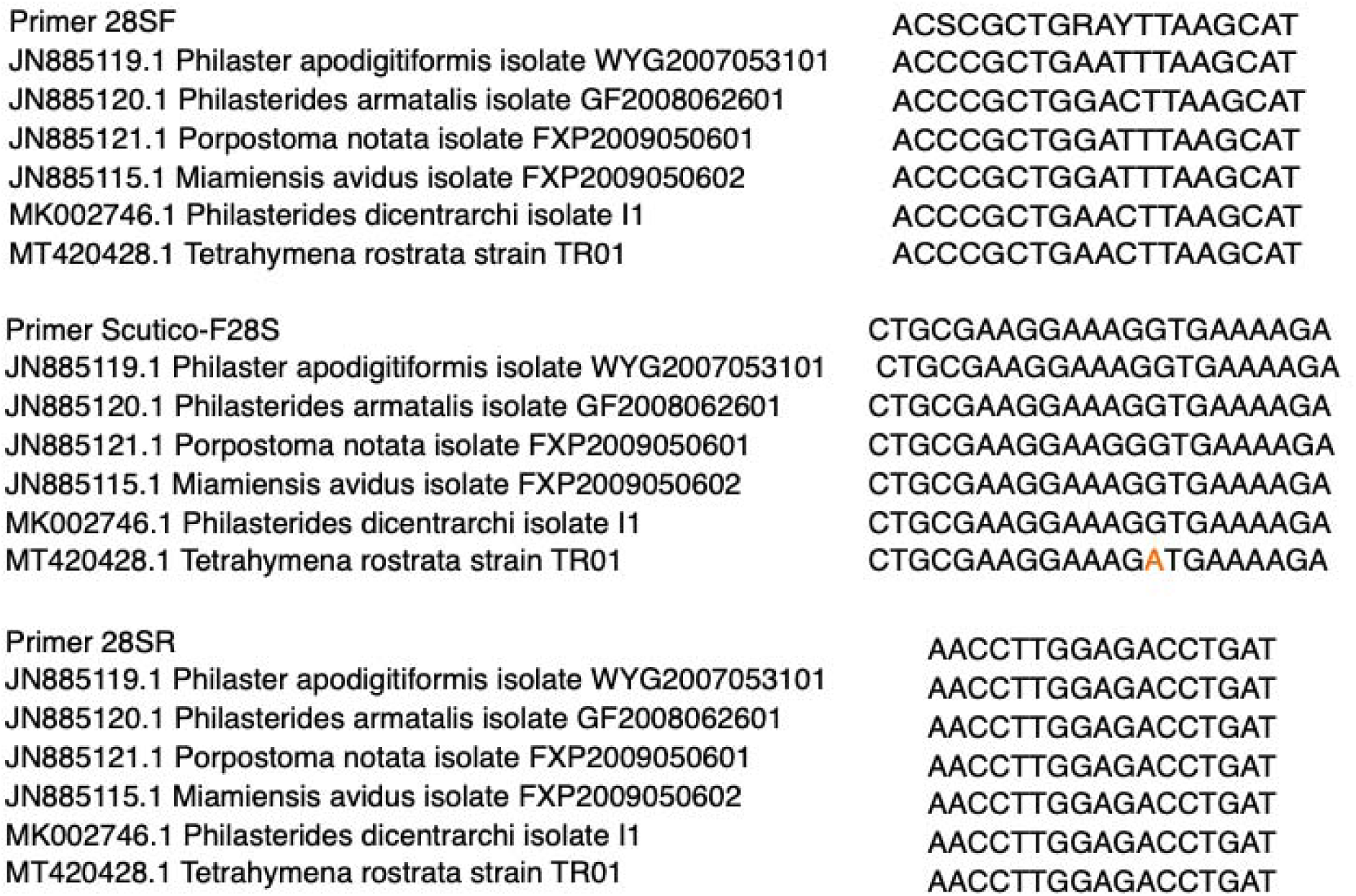
Alignment of primer sequences for 28S rRNA amplification. The figure shows alignment of the primer sequence and region of homology within the 28S rRNA of closely related ciliates. In orange are moderate impact mismatches.

**Supplementary Figure 2.**
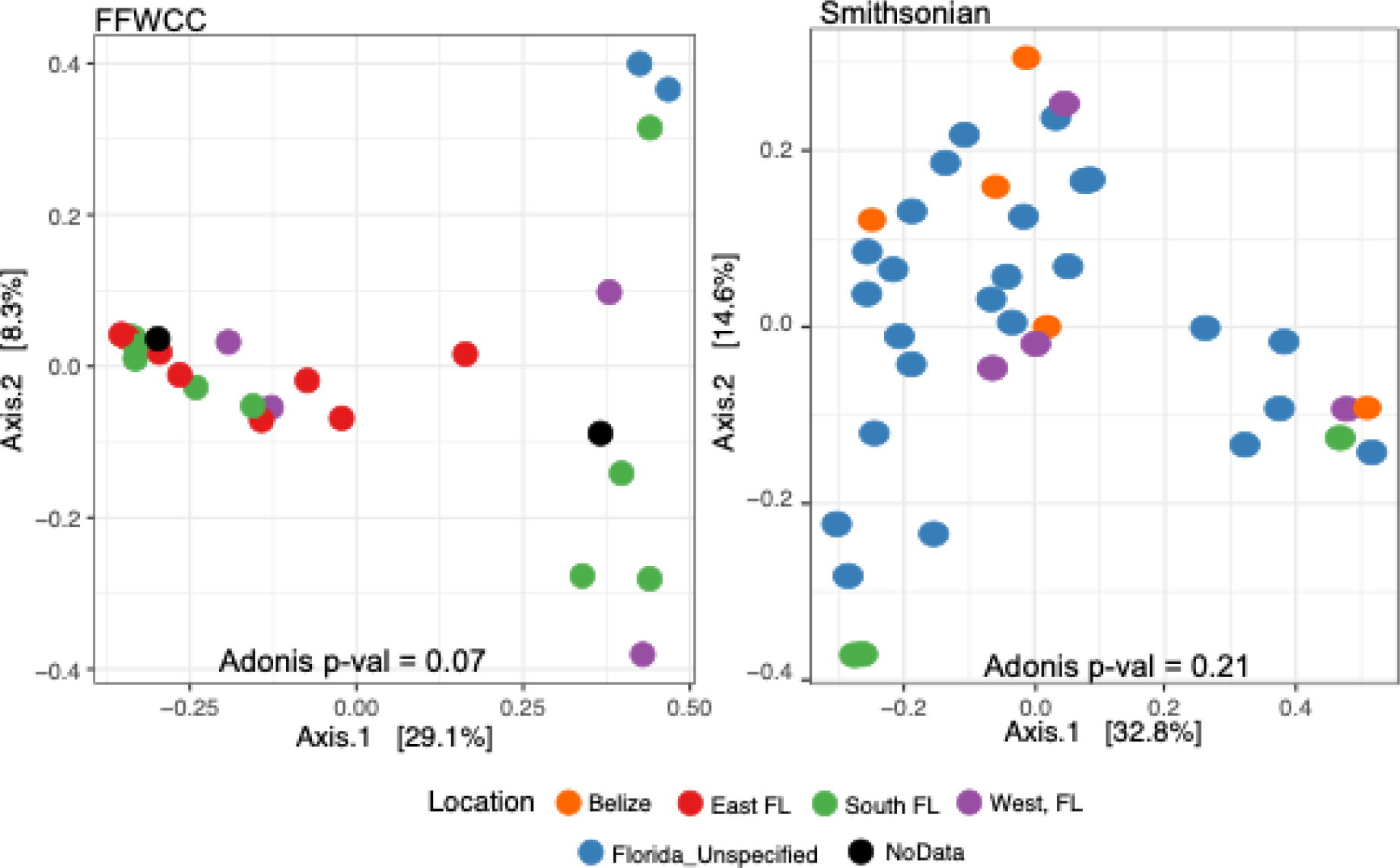
Bray Curtis dissimilarities in a PCoA visualization showing dissimilarity between samples based on collection location divided by storage facility - Florida Fish and Wildlife Conservation Commission (A) or National History Museums Smithsonian Institute (B).

